# Collateral damage: Identification and characterisation of spontaneous mutations causing deafness from a targeted knockout programme

**DOI:** 10.1101/2021.06.30.450312

**Authors:** Morag A. Lewis, Neil J. Ingham, Jing Chen, Selina Pearson, Francesca Di Domenico, Sohinder Rekhi, Rochelle Allen, Matthew Drake, Annelore Willaert, Victoria Rook, Johanna Pass, Thomas Keane, David Adams, Abigail S. Tucker, Jacqueline K. White, Karen P. Steel

## Abstract

Mice carrying targeted mutations are important for investigating gene function and the role of genes in disease, but the process of culturing embryonic stem cells during the making of a targeted allele offers opportunities for spontaneous mutations to arise. Identifying spontaneous mutations relies on the detection of phenotypes segregating independently of targeted alleles, and many phenotypes are easy to miss if not specifically looked for. Here we present data from a large, targeted knockout programme in which mice were analysed through a phenotyping pipeline. Twenty-five lines out of 1311 displayed different deafness phenotypes that did not segregate with the targeted allele. We have identified 8 different mutations causing deafness in 16 of these 25 lines and characterised the resulting phenotypes. Our data show that spontaneous mutations with observable effects on phenotype are a common side effect of intensive breeding programmes, including those underlying targeted mutation programmes.

## Introduction

Manipulating embryonic stem (ES) cells to insert or alter DNA to study the effect of targeted genes in the resulting organism is a widely practised technique, and the creation and characterisation of mutant organisms is a key step in exploring gene function. The advent of the CRISPR-Cas9 system has prompted exploration of its potential unwanted off-target mutagenic effects, which are of particular concern for its use as a therapeutic tool. While some studies have reported that off-target effects are “minimal and manageable”^1, 2^, with no increase in mutation frequency^2, 3^, others have shown that off-target mutations do occur and even alleles present at a low frequency in a G0 mosaic founder could be transmitted to offspring in mice^4^. The creation of targeted mutant alleles involves manipulation of ES cells, and although the mutation frequency is lower in ES cells than in somatic cells^5^, mutations do still occur, particularly when the ES cells have been through multiple passages^6^. Spontaneous mutations may thus arise at any point during the process of making a specific allele, including breeding of the resultant organisms.

The Sanger Institute Mouse Genetics Project was a large-scale programme that generated and screened mice carrying knockdown alleles created by the KOMP (Knock Out Mouse Programme) and EUCOMM (European Conditional Mouse Mutagenesis) programmes^7–9^. Each mouse line established carried a single targeted knockout allele, and animals from each line were put through a wide range of phenotyping tests^9^. One of the tests used was the Auditory Brainstem Response (ABR), a highly sensitive test capable of detecting subtle hearing defects^10, 11^. Vestibular defects leading to balance problems such as circling and head-bobbing were also noted. This enabled the detection of auditory and vestibular phenotypes which did not segregate with the targeted allele and are likely to have been caused by spontaneous mutations. Of the 1311 lines tested, including 2218 wildtype mice and 6798 mutants, we found 25 lines with non-segregating phenotypes.

We observed a variety of phenotypes by ABR and behavioural assessment, and isolated eight lines showing early-onset severe progressive hearing loss, later-onset progressive hearing loss, low frequency hearing loss, or complete deafness with vestibular dysfunction. Here we present our characterisation of these lines, in which we have identified multiple mutations, some of which affect hearing and/or balance. The causative mutations identified so far include new genes not previously associated with deafness and new alleles of genes known to be involved in hearing. Two of the latter show a phenotype much reduced in severity compared to other mutant alleles of the same gene. To the best of our knowledge, this is the first in-depth study of spontaneous mutations in an intensive targeted mutagenesis programme.

## Results

### Identification of spontaneous mutations affecting hearing

From our routine ABR screening of mice carrying targeted knockout alleles, we identified twenty-two lines where a hearing impairment phenotype did not segregate with the targeted allele (Fig. 1, Table 1). An additional four lines were discovered because they displayed a vestibular defect, one of which (MFFD) had also been through the ABR screen, making twenty-five lines in total. Line or colony names (MXXX) were arbitrarily assigned to each breeding colony to use as part of the unique identifier of each mouse and refer to mice carrying both mutant and wildtype alleles within each colony. After each mouse with an aberrant phenotype was discovered, we screened closely-related mice from the same colony and were able to obtain eight mutant lines displaying reliable inheritance of the observed phenotype (Table 1). We used a standard positional cloning approach to identify the mutations. We set up backcrosses to identify the critical chromosomal region for each mutation (Supplementary Fig. 1) and carried out whole exome sequencing to identify candidate mutations within each region. We resequenced candidate variants by Sanger sequencing and tested the segregation of the candidate mutation with the phenotype. For two mutations we confirmed causation using a complementation test. We were able to identify the causative mutation in all eight lines (Table 1), one of which, *S1pr2^stdf^*, has already been described^12^. We then confirmed the presence of one of the mutations in 8 more lines, all displaying the same unusual phenotype. Of the 25 lines with aberrant phenotypes, we have identified the causative mutation in 16 of them.

**Fig. 1.**
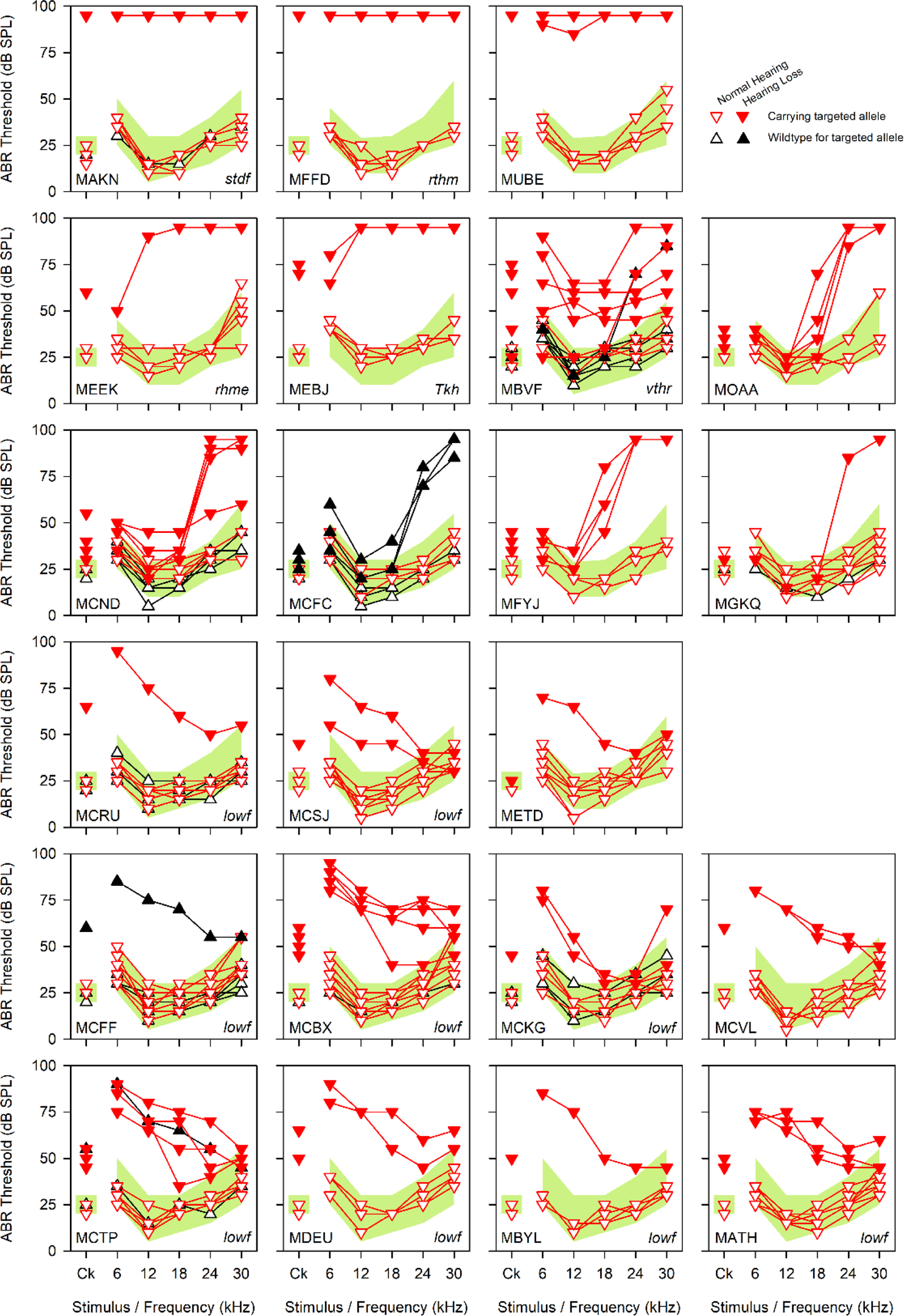
ABR thresholds from 22 targeted lines displaying non-segregating hearing loss. ABR thresholds showing individual mice from targeted mutant lines displaying hearing loss (filled symbols) which does not segregate with the targeted mutant allele (red, inverted triangles) from the original ABR screen. Mice with normal hearing are shown with empty symbols, and black triangles indicate mice which were wildtype for the targeted mutant allele. Affected mice from the MAKN, MFFD and MUBE lines (top) were almost completely deaf. Affected mice from the MEEK and MEBJ colonies exhibited severe hearing loss, while affected mice from the MBVF colony exhibited varying degrees of hearing loss across all frequencies. Affected mice from the MOAA, MCND, MCFC, MFYJ and MGQK colonies (second and third lines) had hearing loss affecting only the higher frequencies, while the remaining eleven colonies all exhibited hearing loss affecting primarily the low frequencies. The shaded pale green area on each panel denotes the 95% reference range for a large population of control wildtype mice derived from littermates of the tested mice, defined in ^11^. All mice were tested at 14 weeks old. Mice from three other lines which exhibited non-segregating hearing loss were detected because of their vestibular phenotype (due to the *Tbx1^ttch^*, *Pcdh15^jigl^* and *Espn^spdz^* alleles), and did not undergo ABRs as part of the ABR screening pipeline. MAKN n=7 unaffected, 3 affected; MFFD n=5 unaffected, 1 affected; MUBE n=4 unaffected, 2 affected; MEEK n=5 unaffected, 1 affected; MEBJ n=4 unaffected, 2 affected; MBVF n=11 unaffected, 6 affected; MOAA n=3 unaffected, 6 affected; MCND n=9 unaffected, 5 affected; MCFC n=7 unaffected, 3 affected; MFYJ n=3 unaffected, 4 affected; MGKQ n=9 unaffected, 1 affected; MCRU n=11 unaffected, 1 affected; MCSJ n=7 unaffected, 2 affected; METD n=7 unaffected, 1 affected; MDEU n=5 unaffected, 2 affected; MCFF n=16 unaffected, 1 affected; MCBX n=9 unaffected, 5 affected; MCKG n=8 unaffected, 2 affected; MCVL n=7 unaffected, 2 affected; MCTP n=7 unaffected, 5 affected; MBYL n=5 unaffected, 1 affected; MATH n=8 unaffected, 3 affected.

**Table 1.**
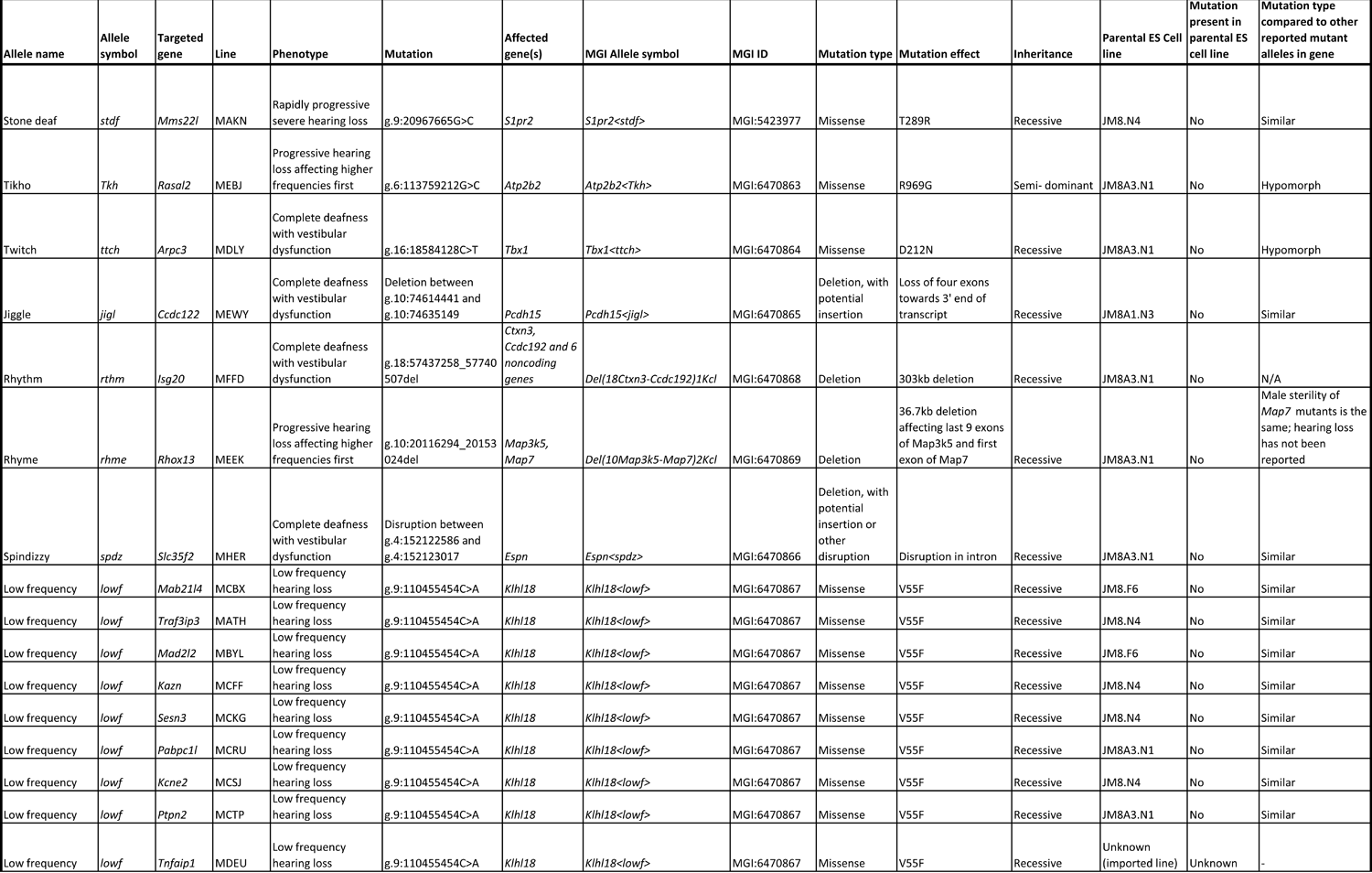
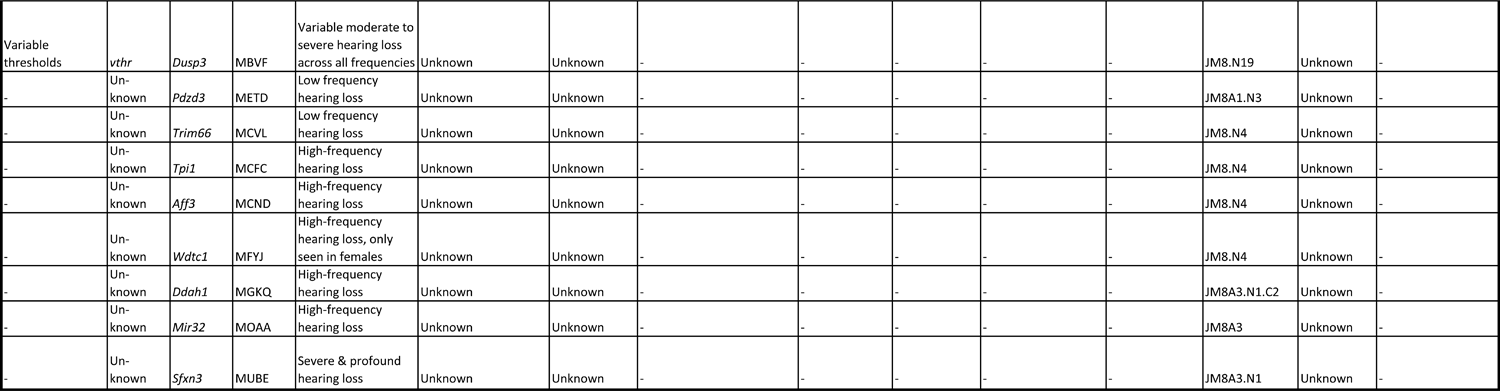
Summary of the 25 lines found to have an spontaneous mutation affecting hearing. This table summarises the 25 lines in which hearing phenotypes were observed which did not segregate with the targeted allele. While some of the targeted genes have been associated with a hearing phenotype (eg *Sesn3* homozygotes have abnormal waveforms^11^, and *ISG20* has been associated with human hearing loss via GWAS^80, 81^), the targeted alleles did not segregate with the phenotypes reported in this paper, and are not linked to the observed hearing loss described here.

### The *Klhl18^lowf^* allele (MCBX colony): low frequency progressive hearing loss

Mice homozygous for this mutation displayed the unusual phenotype of low frequency hearing loss (*lowf*) (Fig. 2a). We mapped the mutation to a 5.7Mb region on chromosome 9 (Supplementary Fig. 1), in which we found 3 exonic variants (Supplementary Table 1), two of which proved to be false calls when resequenced using Sanger sequencing. The third variant was a missense variant in the gene *Klhl18*, g.9:110455454C>A, causing an amino acid change of p.(Val55Phe) (ENSMUST00000068025) (Fig. 2b, c). We used Phyre2^13^ to create a model of KLHL18 based on two structures, a *Plasmodium falciparum* Kelch protein^14^ and the crystal structure of the BTB-BACK domains of human KLHL11^15^ (Fig. 2d). The affected residue lies in the BTB domain, which is involved in protein-protein interaction, including homodimerisation^16–18^. The mutant phenylalanine, being much larger than the wildtype valine, could potentially disrupt the BTB domain and thus protein function. We confirmed that this was the causative mutation by complementation testing with mice carrying the *Klhl18^tm1a(KOMP)Wtsi^* targeted allele^9, 11^ (Fig. 2e). Middle ear dissection and inner ear clearing showed no gross malformations of the ossicles or the inner ear (Supplementary Figs 2, 3). This phenotype was observed in eleven different lines in total (Fig. 1), so we sequenced affected mice from eight more of these lines (in addition to the MCBX line) and discovered that all the affected mice (n=15) were homozygous for the *Klhl18^lowf^* allele, while unaffected littermates (n=8) were heterozygous or wildtype. We were not able to obtain DNA samples from affected mice of the remaining two lines (MCVL, METD). Further characterisation of the unusual *Klhl18^lowf^* phenotype is described in ^19^.

**Fig. 2.**
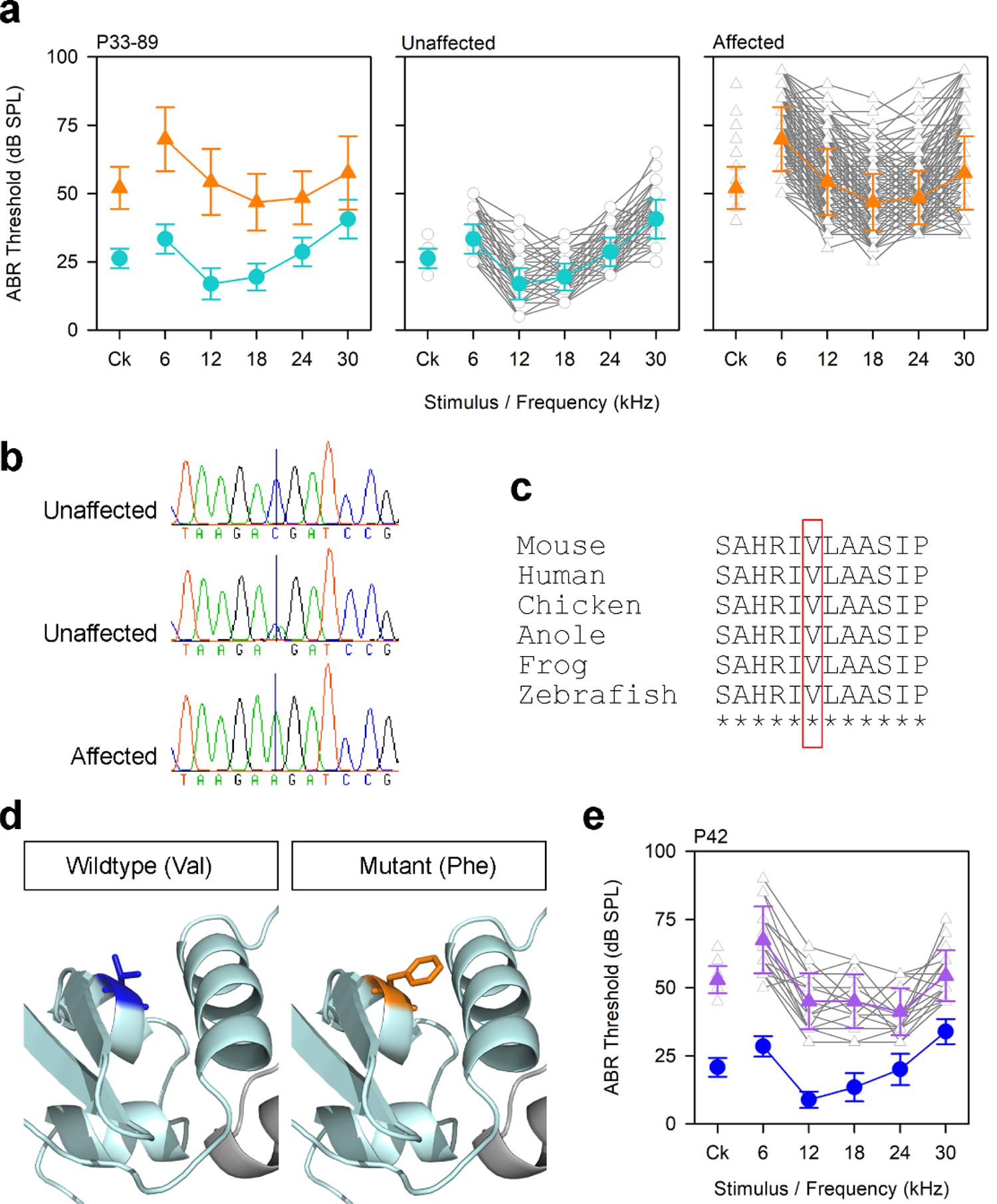
Mutations in *Klhl18* cause low frequency hearing loss. **a** ABR thresholds from mice tested during establishment of the breeding colony derived from the MCBX colony. Mice were tested between 33 and 89 days old, and grouped into affected (n=221, orange triangles) and unaffected (n=213, teal circles) based on the bimodal distributions of thresholds for clicks and 6kHz stimuli. Plots of individual ABR thresholds (grey) are shown separately with the mean trace indicated by coloured lines and symbols; error bars on mean trace are standard deviations. **b** Sequence traces from two unaffected mice (one wildtype, one heterozygote) and an affected mouse (homozygote) showing the variant *Klhl18^lowf^* (MCBX colony), p.V55F. **c** Clustal alignment from mouse, human, chicken, anole lizard, frog and zebrafish showing that the affected amino acid is highly conserved (red box). **d** Close-up of the BTB domain (pale cyan) showing the amino acid structures for the wildtype residue (blue, left) and the mutant residue (orange, right). **e** ABR thresholds from mice heterozygous for the *lowf* allele (n=13, blue circles) and compound heterozygotes carrying the *lowf* allele and the *Klhl18^tm1a^* allele (n=17, purple triangles), demonstrating the low frequency hearing loss phenotype. Individual traces for the compound heterozygotes are shown in grey; error bars on mean trace are standard deviations.

**Fig. 3.**
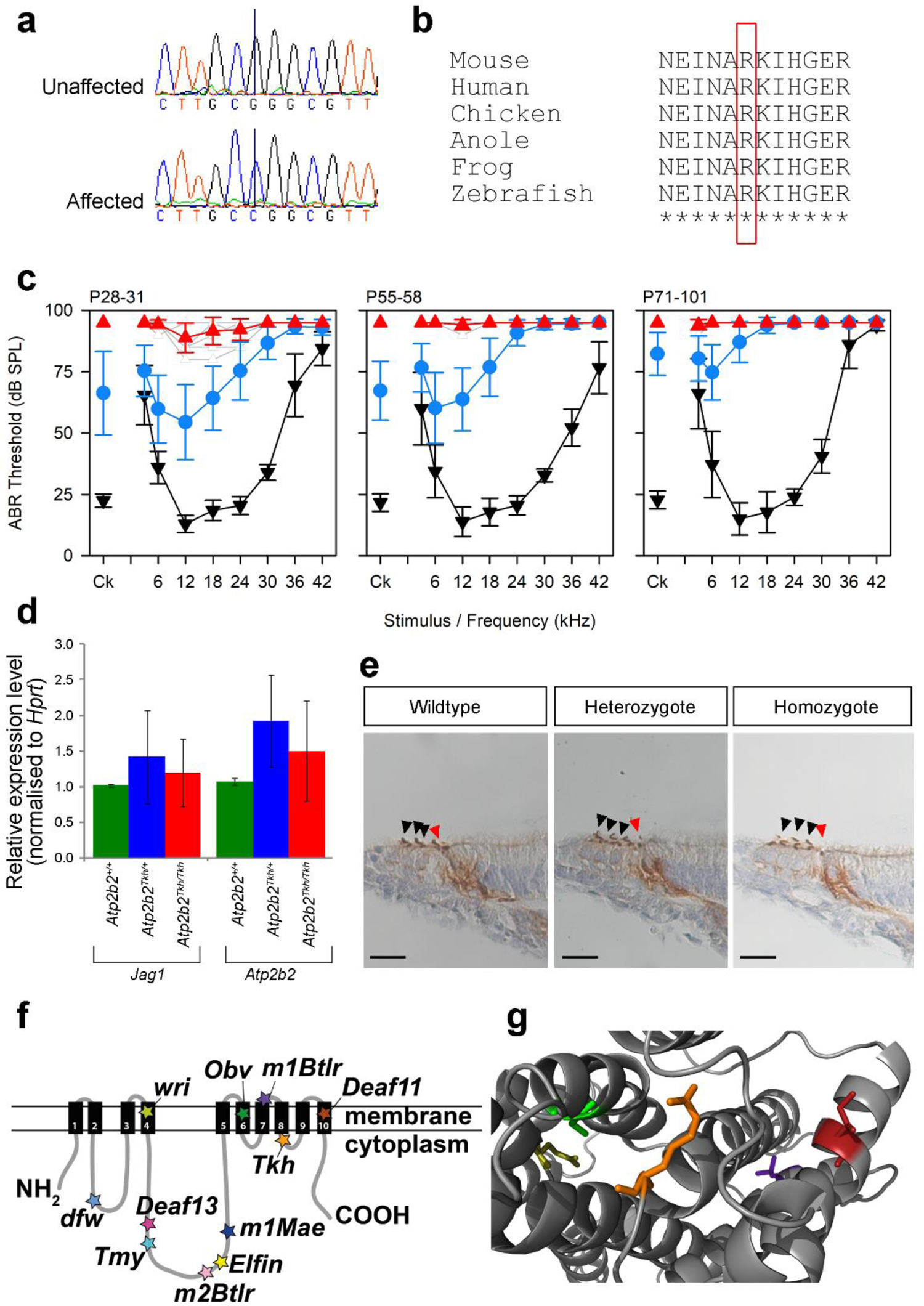
A missense mutation in *Atp2b2* results in semidominant progressive hearing loss. **a** Sequence traces from an unaffected and an affected mouse showing the variant *Atp2b2^Tkh^* (MEBJ colony), p.R969G. **b** Clustal alignment from mouse, human, chicken, anole lizard, frog and zebrafish showing that the affected amino acid is highly conserved (red box). **c** ABR thresholds from wildtype (black inverted triangles), heterozygote (blue circles) and homozygote (red triangles) mice at P28-P31 (n=10 wildtypes, 27 heterozygotes, 9 homozygotes), P55-P58 (n= 9 wildtypes, 24 heterozygotes, 6 homozygotes) and P71-P101 (n=9 wildtypes, 24 heterozygotes, 6 homozygotes). **d** *Jag1* and *Atp2b2* qPCR on RNA from the organ of Corti at P4 (n = 6 wildtypes, 6 heterozygotes and 6 homozygotes). No significant difference was seen between wildtypes, heterozygotes and homozygotes (*Jag1* p=0.357, *Atp2b2* p=0.057, one way ANOVA). **e** PMCA2 antibody stains at P4 (n=2 wildtypes, 2 heterozygotes, 2 homozygotes), showing hair cells from the region 43% of the distance along the organ of Corti from base to apex. Images are representative examples for each genotype, and no differences were observed between genotypes. Arrowheads indicate the hair cells, red for the inner hair cell and black for the outer hair cells. Scale bar = 20µm. **f** Schematic of PMCA2 protein showing the 10 transmembrane helices and the 11 known missense mutations^21–27^ ((https://mutagenetix.utsouthwestern.edu/phenotypic/phenotypic_rec.cfm?pk=1638) and (https://mutagenetix.utsouthwestern.edu/phenotypic/phenotypic_rec.cfm?pk=3670)). **g** Model of PMCA2 protein with the amino acid affected by the *Tkh* allele in orange. The four other missense mutations located in or near the transmembrane helices are also visible, shown in green (*Obv* ^27^), dark red (*Deaf11* ^21^), olive yellow (*wri* ^22^) and purple (*M1Btlr*, https://mutagenetix.utsouthwestern.edu/phenotypic/phenotypic_rec.cfm?pk=1638). The wildtype amino acid structures are shown for all four residues.

### The *Atp2b2^Tkh^* allele (MEBJ colony): semidominant progressive hearing loss

The Tikho (*Tkh*, Russian for “quiet”) mutation was mapped to a 5.2Mb region on chromosome 6 (Supplementary Fig. 1). We found one exonic SNV in this region (Supplementary Table 1), a missense mutation in the *Atp2b2* gene, g.6:113759212G>C, causing an amino acid change of p.(Arg969Gly) (ENSMUST00000101045) (Fig. 3a, b). Mice homozygous for this allele exhibited rapidly progressive hearing loss, while heterozygotes displayed slower progressive hearing loss, with the high frequencies affected first (Fig. 3c, Supplementary Fig. 4). Homozygotes and heterozygotes displayed normal gait and balance. We sequenced mice from the breeding colony and confirmed the segregation of the allele with the phenotypes observed. No gross malformations of the ossicles or inner ear were observed (Supplementary Figs 2, 3). We used qPCR to test RNA expression and immunohistochemistry to study protein localisation, but found no difference between wildtypes, heterozygotes and homozygotes in either test (Fig. 3d, e), suggesting that the variant does not affect mRNA transcription or protein localisation. We modelled the mutation using a model of the rabbit skeletal muscle Ca^2+^-ATPase^20^, which matches 75% of the residues with 100% confidence. This includes the mutant residue, which lies in the cytoplasmic domain between transmembrane domains 8 and 9. There are ten point mutations resulting in amino acid substitutions reported for *Atp2b2* in the mouse (http://www.informatics.jax.org/allele/summary?markerId=MGI:105368, accessed November 2020), all of which result in deafness (*Deaf11*, *Deaf13*^21^, *wri*^22^, *tmy*^23^, *m1Mae*^24^, *Elfin*^25^, *dfw*^26^, *Obv*^27^, *m1Btlr* (https://mutagenetix.utsouthwestern.edu/phenotypic/phenotypic_rec.cfm?pk=1638) and *m2Btlr* (https://mutagenetix.utsouthwestern.edu/phenotypic/phenotypic_rec.cfm?pk=3670)). The three closest are *Obv*, *m1Btlr* and *Deaf11*, all of which lie within 100 amino acids of the *Tkh* residue (Fig. 3f, g).

### The *Del(10Map3k5-Map7)2Kcl* allele (MEEK colony): progressive hearing loss and male sterility

The rhyme (*rhme*) allele was observed to cause male sterility; no pregnancies or pups were obtained from twelve different affected males, paired in matings for at least 66 days. Affected females were fully fertile. We mapped the mutation to a 7MB region on chromosome 10 (Supplementary Fig. 1), but there were no exonic variants in the region, only an interchromosomal translocation from chr10 to chr13 predicted by BreakDancer (Supplementary Table 1), which proved to be a false call when we carried out Sanger sequencing across the predicted breakpoint. We then looked for deletions of whole exons, using the Integrative Genomics Viewer (IGV) to view all reads aligned to the region, and found a large candidate deletion which we confirmed by segregation testing in phenotyped mice. The causative mutation is a 36.7kb deletion, g.10:20116294_20153024del, which includes the last 9 exons of *Map3k5* and the first exon of *Map7* (Fig. 4a). Mutations in *Map7* have been associated with male sterility^28, 29^. Homozygotes for the *rhme* deletion had raised thresholds at high frequencies at 4 weeks old, and the hearing loss progressed with age (Fig. 4b, Supplementary Fig. 4). No gross malformations of the ossicles or inner ear were observed (Supplementary Figs 2, 3). We investigated the hair cells of affected adults using immunohistochemistry and found the organ of Corti was disrupted towards the basal regions, with loss of outer hair cells and disruption and collapse of the tunnel of Corti (Fig. 4c).

**Fig. 4.**
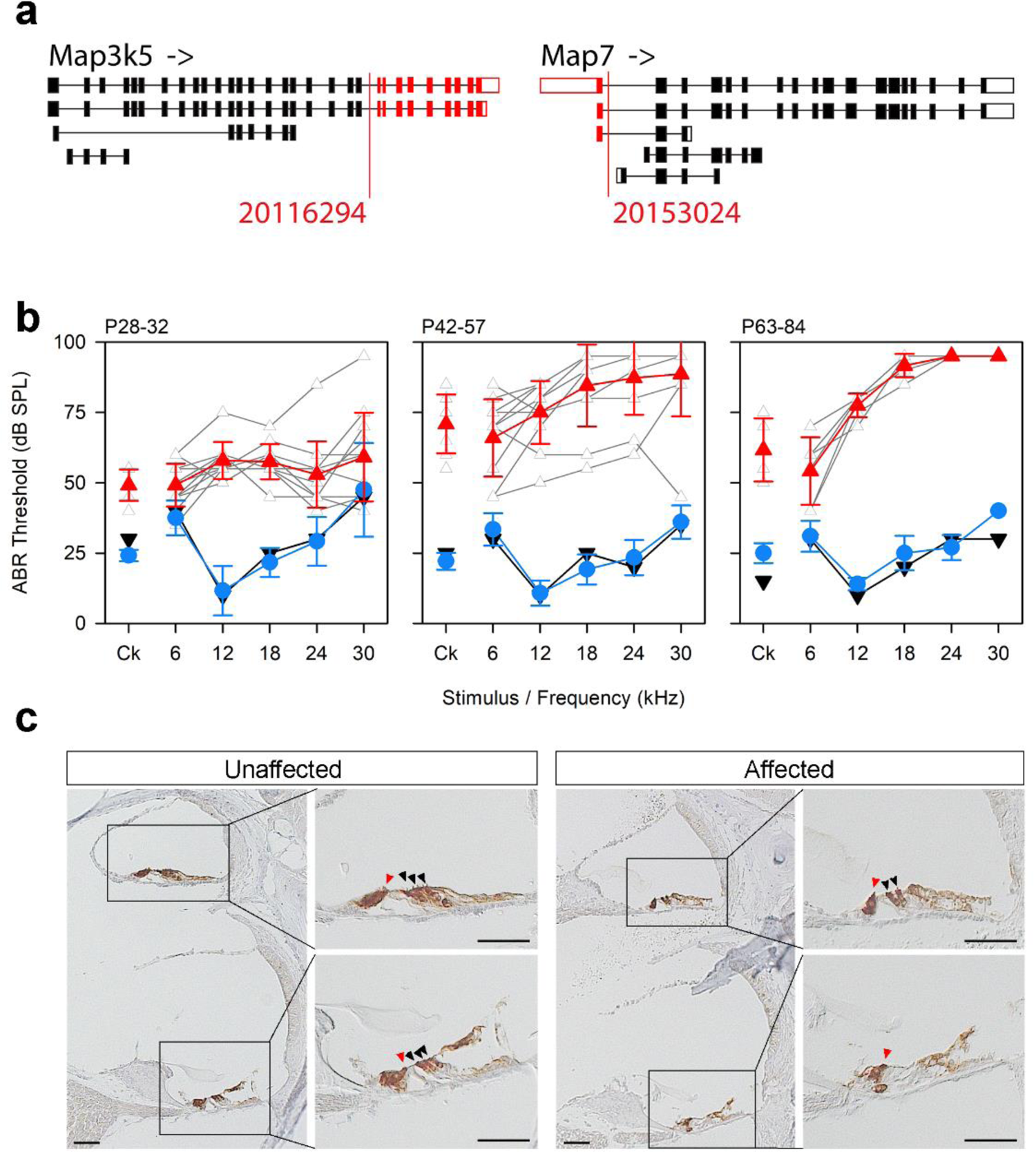
The *rhme* deletion spanning *Map3k5* and *Map7* causes progressive hearing loss and disrupts the organ of Corti. **a** Schematic of the *rhme* deletion seen in the MEEK line showing the different isoforms of the two affected genes (not to scale). Red indicates missing exons. **b** ABR thresholds showing progressive hearing loss in mice homozygous for the deletion (wildtype shown by black inverted triangles, heterozygote by blue circles, homozygotes by red triangles) (n = 1 wildtype, 6 heterozygotes and 12 homozygotes at P28-32; 1 wildtype, 19 heterozygotes and 11 homozygotes at P42-57; 1 wildtype, 5 heterozygotes and 6 homozygotes at P63-84). Traces from individual homozygotes are shown in grey. **c** Expression of MYO7A, a hair cell marker, in the organ of Corti of adult mice showing an apical turn (top; 94% of the distance along the organ of Corti from base to apex), where the hair cells are still present in affected mice, and a mid-basal turn (bottom, 43% of the distance from base to apex), where only the inner hair cell is identifiable (hair cells are marked with arrowheads; red for inner hair cells, black for outer hair cells). Brown indicates where MYO7A is expressed. Three affected and 3 unaffected littermates (at matched ages between 33 and 84 days old) were examined, and the images shown are representative of our observations. Scale bar = 50µm.

### The *Tbx1^ttch^* allele (MDLY colony): complete deafness with vestibular dysfunction

Mice homozygous for the twitch (*ttch*) allele exhibited circling and head bobbing behaviour and had no response to any stimulus up to 95dB, the maximum sound output of our equipment (Fig. 5a). The ossicles were normal in appearance (Supplementary Fig. 2), but we observed signs of inflammation in the middle ear at postnatal day (P)28 (serous effusion, thickened epithelia and capillary hyperplasia) which were more common in affected mice. The gross morphology of the vestibular region was severely affected, with very thin or absent semicircular canals (Fig. 5g). The inner ears of P4 pups displayed a reduced scala media and a thinner stria vascularis at early postnatal stages, although the hair cells appear to have developed normally (Fig. 5d). The saccule had collapsed, but the utricular lumen remained open (Fig. 5d). At adult stages the scala media was even smaller, with a thin or absent stria vascularis, a collapsed Reissner’s membrane, extensive degeneration of the organ of Corti, and spiral ganglion cell loss (Fig. 5e). Both utriculus and sacculus had collapsed, and the hair cells in the utricle appeared disorganised (Fig. 5e). We mapped the mutation to a 3Mb region on chromosome 16 (Supplementary Fig. 1), which contained only 1 exonic SNV (Supplementary Table 1), which segregated with the phenotype in the colony. This was a missense variant in *Tbx1*, g.16:18584128C>T, which results in an amino acid change of p.(Asp212Asn) (ENSMUST00000232335) (Fig. 5b, c). Asp212 is the same amino acid affected by the recently-reported nmf219 mutation^30^, although the underlying coding change (g.16:18584127T>C), and the resulting amino acid change (Asp212Gly), both differ (Fig. 5f). We carried out a complementation test with the *Tbx1^tm1Bld^* allele^31^ and found that the inner ears of compound heterozygotes were notably smaller than those of littermates carrying one copy of the *ttch* allele and had malformed semicircular canals (Fig. 5f). This phenotype, which resembled that of *Tbx1^ttch^* homozygotes, confirmed that the *ttch* allele is the causative mutation.

**Fig. 5.**
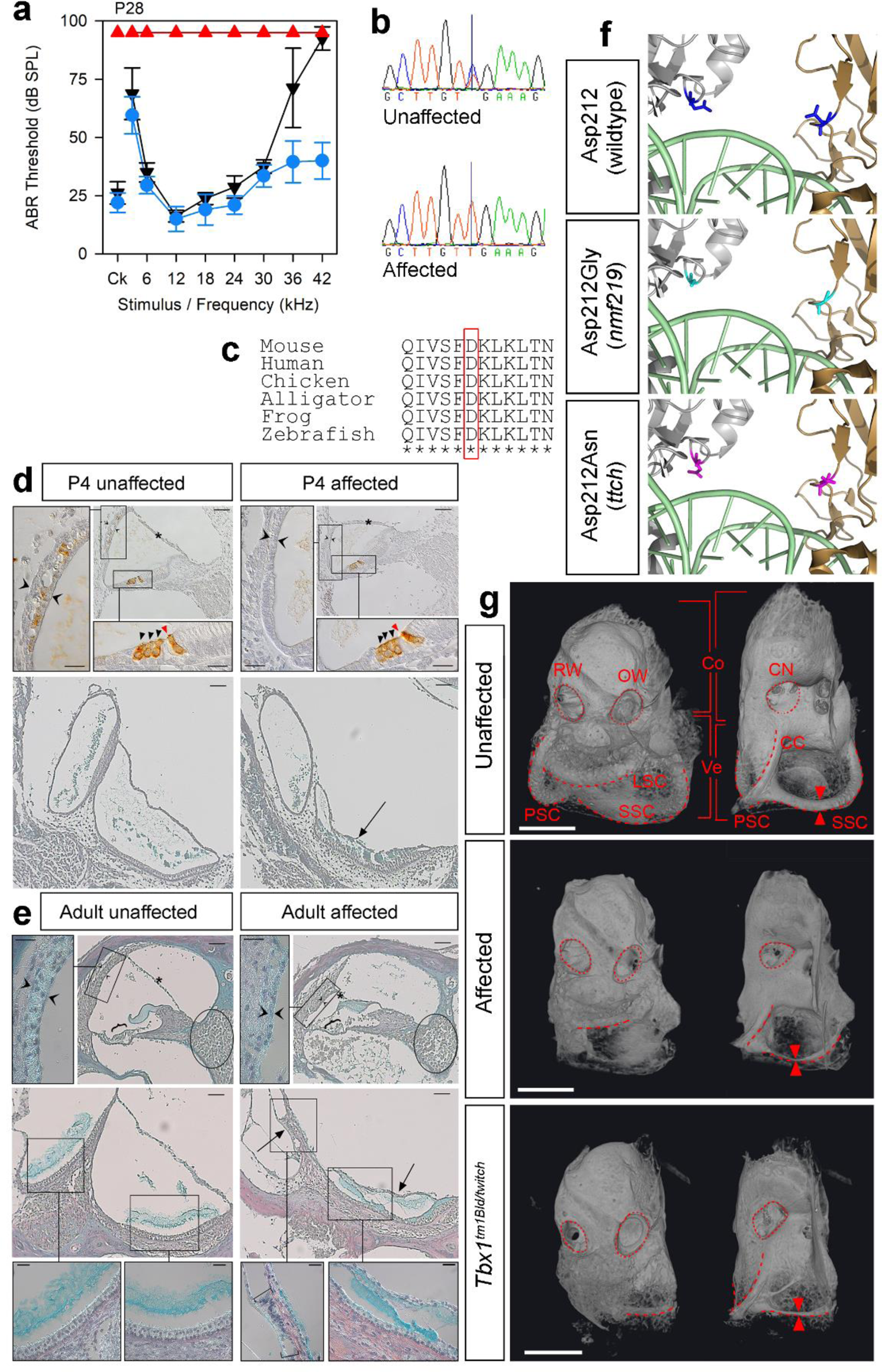
A missense mutation in *Tbx1* causes malformation of the semicircular canals and profound deafness. **a** ABR thresholds from P28 mice homozygous (n=6, red triangles), heterozygous (n=10, blue circles) and wildtype (n=4, black inverted triangles) for the *Tbx1^ttch^* allele (MDLY colony), p.D212N. **b** Sequence traces from an unaffected and an affected mouse showing the variant. **c** Clustal alignment showing conservation of the affected amino acid (red box). **d** P4 sections showing the cochlear duct (top, anti-MYO7A brown stain, blue counterstain, 43% of the distance along the organ of Corti from base to apex) and the maculae (bottom, trichrome staining) (n=3 affected, 3 unaffected littermates). MYO7A is expressed in hair cells (arrowheads; red/black for inner/outer hair cells) and the intermediate cells of the stria vascularis. **e** Trichrome-stained sections from adult mice (n=4 affected, 4 unaffected littermates) showing the cochlear duct (top, 72% of the distance along the organ of Corti from base to apex) and the maculae (bottom). Brackets mark the organ of Corti and the spiral ganglion area is circled. Square brackets indicate abnormal saccular hair cells. In **d** and **e**, main panel scale bars = 50µm, high magnification panel scale bars = 20µm. Asterisks mark Reissner’s membrane, twin open arrowheads the stria vascularis, and arrows the collapsed saccule. **f** Human TBX1 protein model^79^, as a homodimer (silver, gold) bound to DNA (pale green). The Asp212 residue is marked in blue (top), with the *nmf219* mutant residue^30^ in cyan (middle) and the *ttch* mutant residue in magenta (bottom). **g** MicroCT scans of cleared inner ears from affected P28 mice (n=3, middle) and compound heterozygotes (*Tbx1^tm1Bld/ttch^*) at P21 (n=2, bottom). An unaffected P21 mouse is shown at the top (P21 n=4; P28 n=3). Dashed lines outline the semicircular canals, with twin arrowheads for comparison of their width. The middle ear side, with the round (RW) and oval (OW) windows (dotted lines) is on the left, and the brain side, where the cochlear nerve exits (CN, dotted lines), on the right. Brackets indicate the cochlea (Co) and vestibular region (Ve). LSC=lateral semicircular canal; SSC=superior semicircular canal; PSC=posterior semicircular canal, CC=common crus. Scale bar = 1mm.

We also identified two other variants in this line before the non-recombinant region was fully defined; a 27bp inframe deletion in the *Kmt2d* gene which underlies Kabuki syndrome in humans^32^ (Supplementary Fig. 5a), and a missense mutation in the gene *Muc13* (Supplementary Fig. 5c, d). These mutations were confirmed to be present in multiple mice from the MDLY colony but did not segregate with the *ttch* phenotype, and follow-up ABR tests found no effect on hearing in homozygotes of either mutation (Supplementary Fig. 5b, e, g). We investigated the expression of MUC13 in the cochlea at P4 and found it was expressed in hair cells, pillar cells and the basal cells of the stria vascularis (Supplementary Fig. 5f).

### The *Pcdh15^jigl^* allele (MEWY colony): complete deafness with vestibular dysfunction

The jiggle (*jigl)* allele was mapped to a 26.9Mb region on chromosome 10 (Supplementary Fig. 1) in which we found no small exonic variants, but one potential interchromosomal translocation within the gene *Eef2* was identified by BreakDancer (Supplementary Table 1). We investigated this using IGV and found that it was based on a small percentage of the reads covering the region, most of which were correctly mapped, suggesting it was a false call. We then examined the entire non-recombinant region and identified a deletion towards the 3’ end of *Pcdh15*, which includes up to 6 coding exons depending on the transcript. There are 29 protein-coding isoforms of *Pcdh15* (ensembl.org, accessed July 2021), 22 of which contain the affected exons. The deletion results in the loss of the 3’ end of the coding sequence for eleven of those transcripts, and in the loss of internal exons for the remaining eleven (Fig. 6a). We localised the 5’ breakpoint to the region between 10:74614441 and 10:74619826, and the 3’ breakpoint to the region between 10:74634994 and 10:74635149. We used primers designed to amplify a region within the deletion (Supplementary Table 3) to confirm it was not present in 20 affected animals, with 20 unaffected mice from the MEWY colony as controls. *Pcdh15^jigl^* homozygotes display circling and head bobbing and have no auditory brainstem response to any stimulus up to 95dB (Fig. 6b). There were no gross malformations of the ossicles or inner ear (Supplementary Figs 2, 3). Scanning electron microscopy of the organ of Corti at P30 showed that affected mice exhibited hair bundle disorganisation that was most marked in the outer hair cells (Fig. 6c). We observed a similar phenotype in the P5 organ of Corti, with disorganisation of the stereocilia within hair cell bundles, a distorted bundle shape overall, and disoriented bundles (Fig. 6c, d). Hair bundles in the P5 vestibular maculae also lacked the typical staircase organisation (Fig. 6d). These hair bundle defects are similar to those described in other *Pcdh15* mutants and reflect the role of PCDH15 as a component of the tip links between adjacent stereocilia^33, 34^.

**Fig. 6.**
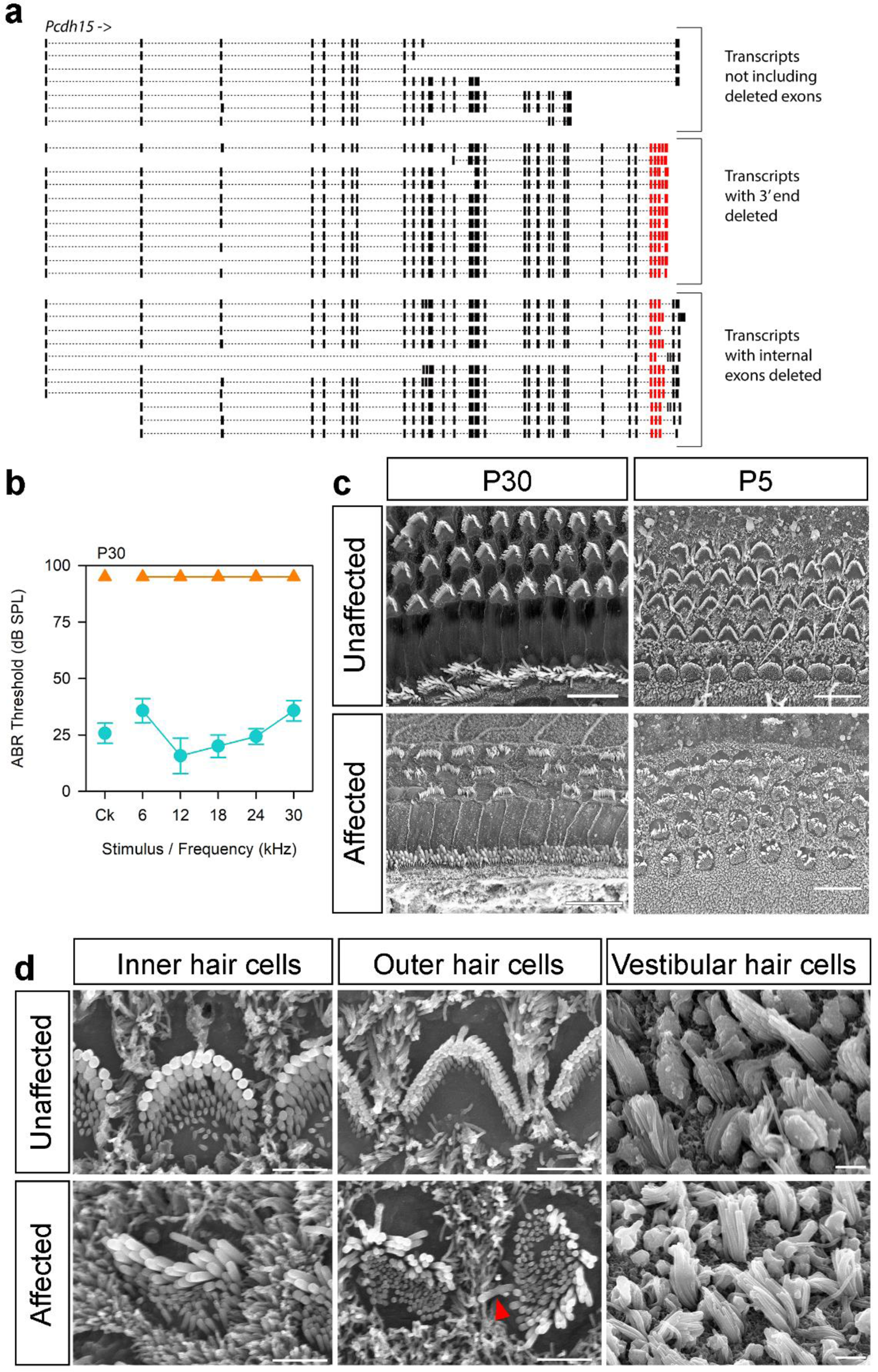
A deletion within the *Pcdh15* gene causes disruption of stereocilia bundles and profound deafness. **a** Schematic of *Pcdh15* isoforms showing the internal deletion in *Pcdh15* (*jigl*, MEWY colony), showing the effects of the deletion on different transcripts (not to scale). Red indicates the deleted exons. Transcripts are shown in their entirety, but not all exons are visible due to scale. **b** ABR thresholds of affected (n=6, orange triangles) and unaffected (n=7, teal circles) mice at P30. **C** Scanning electron micrographs of the organ of Corti at P30 (60-70% of the distance from base to apex, scale bar = 10µm, n=3 unaffected, 6 affected mice) and P5 (50-80% of the distance from base to apex, scale bar = 10µm, n=5 unaffected, 4 affected mice). Mice at P5 were not able to be phenotyped by circling behaviour, but the scanning electron micrographs showed a clear bimodal distribution of affected and unaffected based on disorganisation of hair bundles, and we were unable to amplify the deleted region in affected pups (see Table S7 for primers). Representative examples are shown here. Affected P5 mice had less well-organised hair cell bundles than unaffected littermates. The overall “V” shape of the stereocilia bundle was distorted and irregular in the organ of Corti of affected mice. The polarity of some outer hair cell bundles was rotated by up to 90°, and in some cases the kinocilium was on the opposite side of the cell (arrowhead in **d**). **d** Close ups of hair cells at P5, showing cochlear hair cells from the same region as in (c), and vestibular hair cells from the macula (n= 2 unaffected and 2 affected mice). The arrowhead indicates an example of a kinocilium on the opposite side of the outer hair cell from the stereocilia bundle. Macular stereocilia bundles seemed to have a more ordered staircase structure in the mice classed as unaffected compared to affected mice, where all stereocilia appeared to be long with little sign of a staircase arrangement. Scale bars = 2µm.

### The *Del(18Ctxn3-Ccdc192)1Kcl* allele (MFFD colony): complete deafness with vestibular dysfunction

Mice homozygous for the rhythm (*rthm*) allele exhibited complete deafness (Fig 7a) with circling and head bobbing, suggesting vestibular dysfunction. We mapped the mutation to a 3.3Mb region on chromosome 18 (Supplementary Fig. 1) but did not identify any exonic variants in the region (Supplementary Table 1). We used IGV to examine the nonrecombinant region and discovered a 303kb deletion on chromosome 18, g.18:57437258_57740507del, covering eight genes, including two protein-coding genes (*Ctxn3*, *Ccdc192*), four lncRNA genes, one miRNA and one snRNA (Fig. 7c). This deletion segregated with the phenotype within the colony. The ossicles appeared normal (Supplementary Fig. 2) but the lateral semicircular canal was thinner in homozygotes than in heterozygotes (Fig. 7d), and Myo7a staining revealed severe disruption of the cochlear duct (Fig. 7e). This phenotype resembles that of mice mutant for *Slc12a2*^35^, which lies 138kbp 3’ of the deletion, so we extracted RNA from brain tissue from affected mice and their unaffected littermates at four weeks old, and carried out qPCR to determine whether *Slc12a2* was misregulated. However, there were no significant differences in the levels of *Slc12a2* expression in homozygotes compared to heterozygote littermates (Fig. 7b). Thus, we found no evidence of a position effect of the deletion on *Slc12a2* expression in brain, suggesting it is more likely that a gene within the deletion is responsible for the phenotype.

**Fig. 7.**
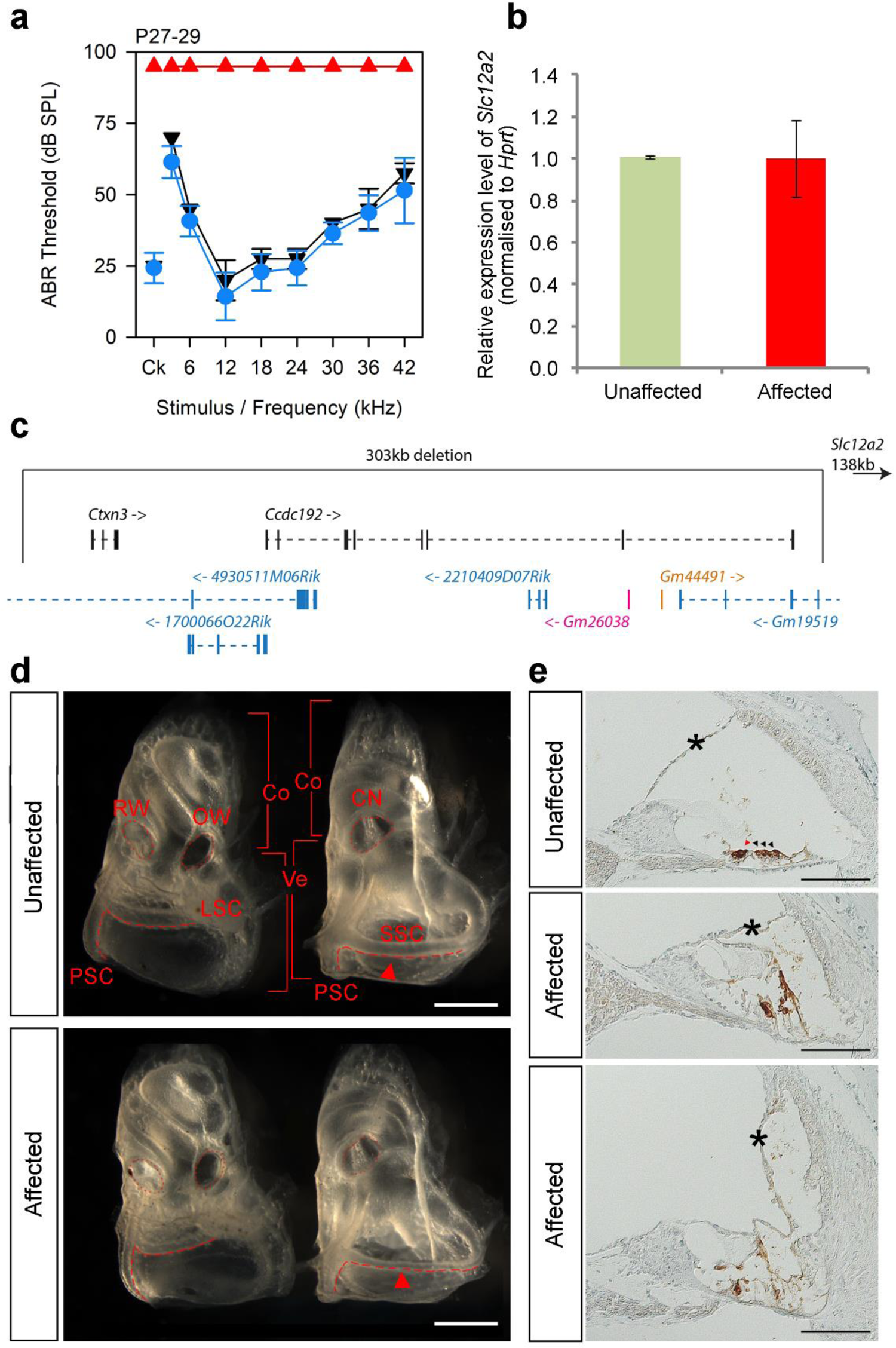
A 303kb deletion on chr18 affects semicircular canals and results in profound deafness. **a** ABR thresholds of mice homozygous (n=6, red triangles), heterozygous (n=7, blue circles) and wildtype (n=2, black inverted triangles) for the *rthm* allele (MFFD colony) at P28±1 day. **b** *Slc12a2* qPCR on RNA from the brains of P28 affected (n=4, red, right) and unaffected (n=4, green, left) mice. There is no significant difference between wildtypes and homozygotes (p=1, Wilcoxon rank sum test). **c** Schematic showing the genes affected by the deletion. There are two protein-coding genes (black), one miRNA gene (orange), one snRNA gene (pink) and four lncRNA genes (blue). *Slc12a2* is located 138kb downstream of the 3’ end of the deletion (indicated by arrow top right). **d** Inner ears from affected (n=3) and unaffected (n=3) mice at P28±1 day. The middle ear side, with the round (RW) and oval (OW) windows is shown on the left of each panel, and the brain side, where the cochlear nerve exits (CN), on the right. The round and oval windows and the cochlear nerve exits are marked by dotted lines, and the semicircular canals by dashed lines. Brackets indicate the cochlea (Co) and vestibular region (Ve). LSC=lateral semicircular canal; SSC=superior semicircular canal; PSC=posterior semicircular canal. Scale bar = 1mm.The superior semicircular canals are thinner in affected mice (red arrowheads). **e** Immunohistochemistry of the cochlear duct of affected (n=3) and unaffected (n=3) mice at P28. Brown indicates the presence of MYO7A, which marks hair cells and the intermediate cells of the stria vascularis. Hair cells are only clearly visible in the unaffected mouse, indicated by the arrowheads (red for the inner hair cell and black for the outer hair cells). Asterisks show the Reissner’s membrane, which is displaced in affected mice. The top two panels show the apical region, 72% of the distance from the base to the apex. The lower panel shows the basal region, at 16% of the base-apex distance. Scale bar = 100mm.

### The *Espn^spdz^* allele (MHER colony): complete deafness with vestibular dysfunction

Mice homozygous for this mutation displayed circling and head bobbing and had no response to sound up to 95dB (Fig. 8a); the allele was named spindizzy (*spdz*). There were no gross malformations of the ossicles and inner ear (Supplementary Figs 2, 3), and MYO7A staining in adults suggested hair cells were present (Fig. 8b). However, scanning electron microscopy showed that there were no stereocilia bundles visible at P28 in homozygotes (Fig. 8c), and at P4, stereocilia bundles were present but disorganised, with thin stereocilia and ectopic stereocilia rows (Fig. 8d). The mutation mapped to a 3Mb region on chromosome 4 (Supplementary Fig. 1) containing 28 protein-coding genes. There were 13 small variants called by Samtools in this region (Supplementary Table 1), none of which were within coding sequence. BreakDancer detected four intrachromosomal rearrangements (Supplementary Table 1), affecting exons of *Klhl21*, *Nol9*, *Acot7*, and an intron of *Plekhg5*, but all four are based on a very low proportion of the reads in each homozygote. The only known deafness gene in the non-recombinant region is *Espn*, mutations in which result in shorter, thinner stereocilia at birth, followed by degeneration of stereocilia in early adulthood^36–38^. We therefore sequenced *Espn* mRNA from the brains of adult affected mice and their unaffected littermates. No splicing errors were observed for most of the exons; however, we were unable to amplify sequence from exons 15 and 16 in homozygotes (Fig. 8e). Exon 15 (ENSMUSE00001290053) is a 12bp exon located 85bp from exon 14, and separated by 2.35kb from exon 16, which is the final exon (Fig. 8e). We resequenced the genomic region between exons 14 and 16 in three *spdz* homozygotes using Sanger sequencing, and while no variants were observed in intron 14-15 or in exon 15, we were unable to amplify a 431bp region in the intron between exons 15 and 16, between g.4:152122586 and g.4:152123017, in any of the homozygotes. We successfully sequenced this region in two wildtype mice. This failure to amplify suggests that there may be an insertion or other genomic disruption at this location of the chromosome. We extracted DNA from 55 affected mice from the colony, and in all of them this specific sequence failed to amplify, while an adjacent sequence worked. In 56 unaffected mice both sequences were amplified (Fig. 8e,f; primers in Supplementary Table 3).

**Fig. 8.**
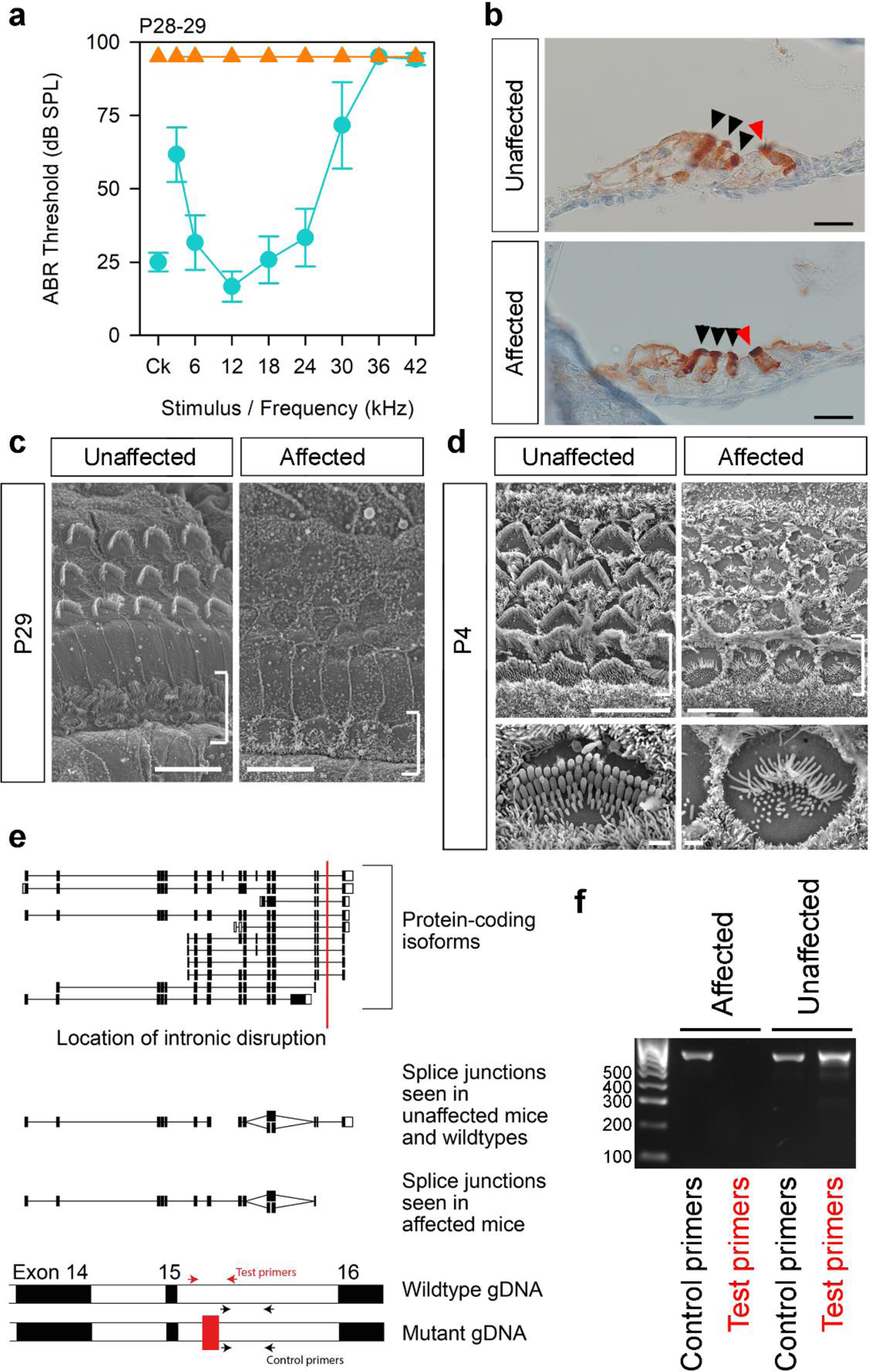
Mice carrying the *spdz* allele (MHER colony) are profoundly deaf, with abnormal hair cell stereocilia at P5 and hair cell degeneration by P29. **a** ABR thresholds of affected (orange triangles, n=7) and unaffected (teal circles, n=6) mice at P28-29. **b** Expression of MYO7A (brown) in hair cells from the apical region of the organ of Corti (90-95% of the distance from base to apex) of mice at P28 (n=3 affected, 3 unaffected), showing that hair cells are visible in affected mice (arrowheads, red for the inner hair cell and black for outer hair cells). Scale bar = 20µm. **c** Scanning electron micrographs of the 12kHz best-frequency region (65-70% of the distance from base to apex) of the organ of Corti in unaffected (n=4) and affected (n=3) mice at P29. Brackets indicate the inner hair cell row. Most hair bundles are missing. Scale bar = 10µm. **d** Scanning electron micrographs of a homozygous mutant mouse at P4 (n=3) and an age-matched wildtype, showing the 42kHz best-frequency region (20% of the distance from base to apex). Brackets indicate the inner hair cell row; scale bar = 10µm. Lower panels show a magnified view of inner hair cells in the same region (scale bar = 1µm). **e** Schematic showing the known protein-coding isoforms of *Espn* and the location of the intronic disruption (red line). The splice junctions identified in brain cDNA are shown below; each line indicates a splice junction visible in at least one mouse. At the bottom is a schematic of the genomic DNA (gDNA) showing exons 14, 15 and the start of exon 16 in black, separated by introns in white (not to scale). The disruption in the mutant is shown by the addition of a red block in the intron. The “test” primer pair (red arrows) fails to amplify in mutant gDNA, and the “control” primer pair (black arrows) works for both wildtype and mutant. **f** Gel showing bands amplified using these primer pairs (Supplementary Table 3) on an affected mouse and an unaffected mouse. The size of ladder bands is shown at the side in bp.

### Three other lines which underwent exome sequencing

In addition to the above seven lines, mice from three other lines displaying nonsegregating phenotypes were sequenced (MAKN, MATH and MBVF colonies, Fig. 1). The mutation underlying the rapidly progressive hearing loss phenotype seen in the MAKN line was a point mutation in the *S1pr2* gene, named stonedeaf (*S1pr2^stdf^*). Mice homozygous for this allele displayed a rapid reduction in endocochlear potential (EP) between P14 and P56 which correlated with the progression of their hearing loss, while hair cell degeneration followed at a later age^12^.

In the case of the mutation in the MATH line we were not able to establish a breeding colony inheriting the phenotype, but we were able to confirm the presence of the *Klhl18^lowf^* mutation from the exome sequence, which is in accordance with the low frequency hearing loss exhibited by these mice (Fig 1).

The affected mice from the MBVF line displayed variably raised thresholds across all frequencies (Fig 1); we named the allele variable thresholds, *vthr*. However, because of this extreme variability, we were unable to carry out a backcross or maintain the *vthr* phenotype within the colony. We did observe that the thresholds of 7 affected mice, while variable at 14 weeks old, progressed to more severe hearing loss at 6 months old (Supplementary Fig 4c). We collected and examined the middle ears of these affected mice and related unaffected mice at ages over 6 months and did not observe any middle ear defects (n=7 affected, 17 unaffected). Because of this variability in ABR thresholds, we did not restrict the zygosity of identified variants during variant processing (Supplementary Table 1). After quality and impact filtering, we found 221 potential high impact variants, including 15 large structural variants that were predicted to affect 30 known deafness genes between them (Supplementary Table 2).

### ES Cell sequencing

The 25 lines with spontaneous mutations were derived from several different parental embryonic stem cell lines (Table 1). We carried out whole exome sequencing on three of these lines (JM8.F6, JM8.N4 and JM8.N19) and used Sanger sequencing of selected regions to check two others (JM8A1.N3 and JM8A3.N1)^39^. For the eight alleles affecting hearing described above (including *S1pr2^stdf^*), we found that the mutations were not present in the parental ES cell lines (Table 1). The *Klhl18^lowf^* mutation, the only one seen in multiple lines, was not found in any of the ES cell lines from which we obtained sequence (Table 1).

In order to find out whether any variants seen in the mice could have been derived from the parental ES cell lines, we compared the JM8.F6 and JM8.N4 whole exome sequencing to whole exome sequencing from four descendant mice (two from the MCBX line and two from the MATH line, both of which carried the spontaneous *Klhl18^lowf^* mutation, Fig 1, Table 1). For this we adapted the variant filtering steps and processed exome sequence data from each mouse independently (Supplementary Table 4a). We compared the high-quality variants identified by the different callers and found that a subset of ES cell variants was indeed found in the mice created from each line (e.g. 8 out of 1105 variants called by SAMtools were found in the JM8.F6 ES cells and in both MCBX mice, Supplementary Fig 6). We chose 21 high quality, high impact variants for confirmation by Sanger sequencing but most variants were not validated in either the ES cells or the mice (Supplementary Table 4b). However, two variants were found in both JM8.F6 and the two MCBX mice; one point mutation identified by SAMtools and one small indel called by Dindel (Supplementary Table 4b). These variants were not seen in the two MATH mice, which shared the same low frequency hearing loss mutation and the spontaneous *Klhl18^lowf^* allele, nor in the JM8.N4 ES cell sequence.

### Potential sources of the spontaneous mutations affecting hearing

As none of the spontaneous mutations we found that affected hearing came from the original parental ES cell lines (prior to genetic manipulation), we examined the pedigrees of the mice originally found to have these phenotypes and the mice used as founders for our eight breeding colonies. For each line, we identified the latest possible point at which the mutant allele could have arisen, assuming that it only occurred once. In five lines, the mutation could have arisen just two generations before it was observed (MEBJ (*Tkh*), MHER (*spdz*), MEEK (*rhme*), MDLY (*ttch*) and MEWY (*jigl*)). In the MAKN (*stdf*) line, the mutation must have arisen no less than three generations before the phenotype was observed (Supplementary Fig 7). However, in the MFFD (*rthm*) line, the mutation must have been present in the chimaeric offspring of the microinjected founder (Supplementary Fig. 7). For the lines bearing the *Klhl18^lowf^* allele, the most likely source is the wildtype colony used for embryo donors, microinjection and colony expansion. We checked the pedigrees of the mice homozygous for the *Klhl18^lowf^* allele and confirmed that in each case there was an ancestral mouse from the same wildtype colony which could have passed on the *Klhl18^lowf^* allele to both the dam and sire of the homozygous mouse. We then constructed pedigrees for these wildtype mice and determined that it is likely that the *Klhl18^lowf^* allele was present in several of the founders of the C57BL/6N wildtype colony used to expand the mutant lines^9^.

In summary, one mutation was present at the point of microinjection (MFFD, *rthm*) and therefore arose when the ES cells were targeted or during the ES cell processing prior to microinjection, and one is likely to have arisen spontaneously in a wildtype colony (MCBX, *Klhl18^lowf^*). The remaining six mutations could have occurred either during ES cell targeting and processing, or during breeding of the colony carrying a targeted allele (Fig 9).

**Fig. 9.**
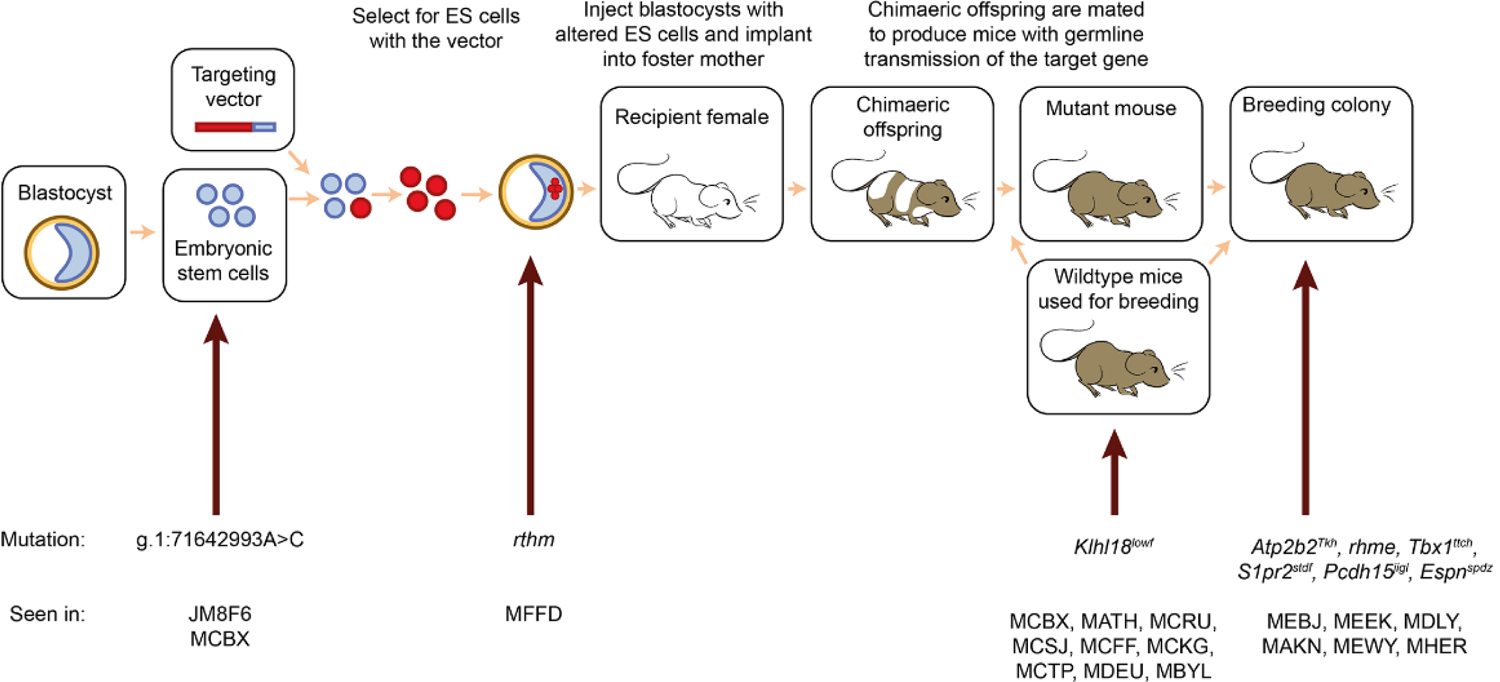
There are many opportunities for spontaneous mutations to arise during the process of making a targeted knockout allele. A schematic showing the stages of making knockout mice. Spontaneous mutations can arise at any point in this process. The mutations described in this paper are shown at the bottom, with arrows indicating the latest possible time at which that mutation could have occurred. If a spontaneous mutation occurs in the cultured embryonic stem cells before targeting, it has the potential to affect multiple mouse lines, but although we detected variants in MCBX mice which were present in the parental ES cell line JM8F6 (eg g.1: 71642993A>C, Supplementary Fig 6, Supplementary Table 4b), none of the eight mutations affecting hearing were found in any of the parental ES cell lines (Table 1). A mutation arising later in the process may be specific to a single line (such as the *rthm* allele, which is likely to have arisen in the targeted ES cell) or a single mating within that line, which is likely to be the case for most of the mutations described here. The *Klhl18^lowf^* mutation, on the other hand, probably arose within the wildtype line used for expansion for all the colonies in which mice with the low frequency hearing loss phenotype were found.

## Discussion

Here we have described seven mutant alleles affecting hearing which, along with the previously described *S1pr2^stdf^* mutation^12^, arose as spontaneous mutations within a targeted knockout programme (Table 1). In total, we observed 25 lines with hearing impairment which did not segregate with the targeted allele (Fig. 1), and we identified the causative mutation in 16 of them. It’s likely that in the cases where we could not establish a breeding colony carrying the phenotype, the related mice we obtained weren’t carrying the mutation. However, it’s possible that in the lines where only one mouse was found to be affected, the mutation causing hearing loss was a somatic mutation (eg MGKQ, Fig. 1). In one case (MBVF), the phenotype was variable, and could not be reliably maintained. We have identified six novel alleles of known deafness genes (*Klhl18* ^10, 11^, *S1pr2* ^12, 40–42^, *Atp2b2*^21–27, 43^, *Tbx1*^44–47^ *Pcdh15*^48^ and *Espn*^49^), and four candidate deafness genes (*Ctxn3*, *Ccdc192*, *Map3k5* and *Map7*). These last lie within the *rthm* and *rhme* deletions in the MFFD and MEEK lines, and more study is required to identify which gene is responsible for the hearing phenotype or whether, in the case of the *rthm* deletion, it is one of the noncoding RNA genes that underlies the deafness and vestibular dysfunction in this line.

Of the known deafness genes, two of our mutations are hypomorphs, *Atp2b2^Tkh^* (MEBJ) and *Tbx1^ttch^* (MDLY) (Table 1). In the case of *Tbx1*, mice homozygous for a null allele die at birth due to failure to inflate their lungs, but gross morphological abnormalities are evident as early as E8.5^44^. mRNA dosage experiments have shown that a very small reduction below the mRNA expression level seen in heterozygotes is sufficient to induce full neonatal lethality^50^. One other *Tbx1* hypomorph has been reported, the *Tbx1^nmf219^* mutant, which carries a missense mutation affecting the same amino acid as the *Tbx1^ttch^* allele, Asp212^30^. *Tbx1^nmf219^* and *Tbx1^ttch^* homozygotes display a similar inner ear phenotype, with hypoplasia of the semicircular canals, a reduced scala media, collapsed Reissner’s membrane, reduced or absent stria vascularis, degeneration of the organ of Corti, and spiral ganglion cell loss^30^. TBX1 binds to DNA as a dimer, and Asp212 is located between the two monomers, so amino acid changes could potentially affect the structure of the dimer and its capacity to bind DNA^30^ (Fig. 5c). It is interesting to note that while many other mutant alleles of *Tbx1* are homozygous perinatal lethal^31, 44, 45^, including at least one point mutation^51^, neither of these Asp212 mutants show reduced viability of homozygotes. *Atp2b2* mutants also display a difference in phenotype depending on the level of activity of PMCA2, the encoded protein^52^. There are 15 reported *Atp2b2* mutants with phenotypes, 10 of which result from a single amino acid change (Fig 3, http://informatics.jax.org, accessed December 2020). Only the *Deaf11* and *Deaf13* missense alleles are like the *Tkh* allele in that they do not result in ataxia in homozygotes^21–27, 43, 52–54^ ((https://mutagenetix.utsouthwestern.edu/phenotypic/phenotypic_rec.cfm?pk=1638) and (https://mutagenetix.utsouthwestern.edu/phenotypic/phenotypic_rec.cfm?pk=3670)). The new hypomorphic alleles reported here may be useful for in-depth studies of the function of these genes.

It is likely that over a thousand genes contribute to the development and function of the ear, making hearing impairment relatively sensitive to background mutation rates^9, 11^. From the 9016 mice (2218 wildtype mice and 6798 targeted mutants) screened by ABR in the current study, 55 had a non-segregating phenotype (0.6%). However, seven of the eight mutations causing a hearing phenotype were inherited in a recessive way, so that we had to screen a mouse with two copies to detect it, and only four mice per colony were normally screened by ABR, although in some cases wildtype littermates were included in the control cohort. Thus, the number of new mutations we found causing deafness is likely to be an underestimate, as many colonies harbouring new mutations causing deafness will not have led to a homozygote reaching the screening cohort. Like many phenotypes, hearing impairment in a mouse, when unaccompanied by vestibular dysfunction, is a subtle phenotype and easy to miss if it is not actively investigated. It is highly likely that multiple other mutations which have no impact on hearing nor any gross impact on the appearance, behaviour or viability of the mice exist within all these lines. We found two such mutations in the process of sequencing the MDLY (*ttch*) mice: a missense mutation in *Muc13* and a deletion in the Kabuki syndrome gene *Kmt2d* (also known as *Mll2*), neither of which affected hearing nor had any obvious effect on homozygotes (Supplementary Fig. 5).

When we first observed the non-segregating phenotypes, particularly the widespread *Klhl18^lowf^* mutation, we suspected that the mutations arose in the ES cells used to make the mice, either before or after insertion of the manipulated allele. Indeed, a previous missense mutation in *Atp2b2* has been reported to have arisen during clonal expansion of targeted ES cells^24^, and multiple other examples of off-target mutations arising in targeted ES cells have been described^55, 56^. However, when we sequenced the parental ES cell lines, we did not find any of these mutations, and from our pedigree analyses, only one can be unambiguously traced to the founder of the line (MFFD, *rthm*), although since we can only identify the latest possible time of occurrence from the pedigrees, we can’t rule out the possibility that the other mutations occurred within the targeted ES cell. In addition, the presence of the *Klhl18^lowf^* mutation in multiple lines suggests that it arose within the wildtype colony used for expansion of the mutant lines (Fig 9). We therefore suggest that it is likely that at least some of the mutations affecting hearing arose during breeding. Thus, regardless of the method of genome manipulation or how long ago a mutation was made, the potential for spontaneous and off-target mutations to affect a mutant animal being studied must always be borne in mind. In particular, a non-segregating mutation that affects the phenotype under study could result in apparently variable penetrance or expressivity of the phenotype. It’s possible that the variable phenotype observed in the MBVF (*vthr*) line (Fig. 1) is the result of more than one spontaneous mutation.

As well as being a reminder to pay attention to unexpected phenotypes, these results highlight the critical importance of proper data collection and retention in any study. The accurate breeding records and careful tracking of the thousands of mice tested in the Mouse Genetics Project were critical to the detection, isolation and identification of these new mutations. Furthermore, confirmation of candidate variants by Sanger sequencing was essential to identify false calls from exome sequence.

## Methods

### Ethics statement

Mouse studies were carried out in accordance with UK Home Office regulations and the UK Animals (Scientific Procedures) Act of 1986 (ASPA) under UK Home Office licences, and the study was approved by both the Wellcome Trust Sanger Institute and the King’s College London Ethical Review Committees. Mice were culled using methods approved under these licences to minimize any possibility of suffering.

### Generation of mice

Mice carrying knockout first conditional-ready alleles were observed for gross behavioural abnormalities using a modified SHIRPA test at 9 weeks old, and tested by ABR at 14 weeks old, as described in ^9^ and ^11^. Mice assigned to the phenotyping pipeline could not be withdrawn for investigation of these non-segregating phenotypes, so mice from the same line, as closely related to the affected mice as possible, were used to set up colonies to screen for the spontaneous mutation. 25 lines were found to carry an spontaneous mutation involving hearing impairment, and eight new breeding colonies were successfully established with the observed phenotype reliably inherited. All mutations were generated and maintained on the C57BL/6N background. These eight mouse lines will be available via the European Mouse Mutant Archive. Two further mouse lines were used for complementation testing of compound heterozygotes: *Klhl18^tm1a(KOMP)Wtsi^* ^9^ and *Tbx1^tm1Bld^* ^31^.

### Auditory Brainstem Response (ABR)

ABR tests were carried out as previously described^11, 57, 58^. We used a broadband click stimulus and shaped tonebursts at a range of pure tone frequencies at sound levels from 0-95dB SPL, in 5dB steps. 256 sweeps were carried out per frequency and sound level, and these were averaged to produce the ABR waveform. A stack of response waveforms was used to identify the threshold for each stimulus, the lowest intensity at which a waveform could be distinguished.

### Assessment of balance

Mouse behaviour was observed in their home cage for signs of circling or headbobbing. To detect balance dysfunction, we also used the contact righting reflex test. Mice were placed in a large petri dish sized such that the back of the mouse was in contact with the dish lid but not under pressure. The petri dish was inverted, and the mouse was monitored for 30 seconds. If it did not turn itself over within that time, it was marked as affected, while if it turned itself over promptly, it was marked as unaffected. Some mice righted themselves after a delay and were tested a second time. If the results of the righting test were unclear, the mouse was not used for mapping.

### Linkage mapping

For each line, affected males were outcrossed to C3HeB/FeJ females. For the lines displaying recessive inheritance, the offspring were backcrossed to affected mice of the same line. For the single line displaying semidominant inheritance (MEBJ, *Atp2b2^Tkh^*), the outcross offspring were backcrossed to C3HeB/FeJ wildtype mice. Backcross offspring were assessed by ABR or contact righting reflex, then after culling, tissue was collected for DNA extraction. The initial genome scan was carried out using a standard marker panel (Supplementary Table 5), then once the linkage region was established, it was narrowed down using more backcross mice, more markers within the region and, where necessary, strain-specific SNVs (Supplementary Table 5).

### Exome sequencing

Two affected mice of each line were selected for exome sequencing. DNA was extracted using phenol and chloroform, and exome sequencing was carried out by the Wellcome Trust Sanger Institute (WTSI), with the exception of the *spdz* sequencing, which was carried out by Novogene (Hong Kong). Genomic DNA (approximately 1 ug) was fragmented to an average size of 150 bp (WTSI) or 180-280bp (Novogene) and subjected to DNA library creation using the Agilent SureSelect Mouse All Exon Kit. Adapter-ligated libraries were amplified and indexed via PCR. Enriched libraries were subjected to 75bp paired-end sequencing (HiSeq 2000; Illumina), with the exception of the *spdz* libraries, which were subjected to 150bp paired-end sequencing (HiSeq 4000; Illumina) following the manufacturer’s instructions. Sequences were aligned to GRCm38 using bwa^59^, with the exception of the *spdz* fastq files, which were checked and processed using FastQC^60^ and Trimmomatic^61^, then aligned to GRCm38 using hisat2^62^. All bam files were improved by local realignment around insertions and deletions discovered in the mouse genomes project^63^ using GATK^64^. Bam files were processed using Picard and Samtools^65, 66^. Raw data can be downloaded from the European Nucleotide Archive, studies PRJEB2585, PRJEB5221 and PRJEB45713. Individual sample accession numbers are in Supplementary Table 6c.

### Variant identification and investigation

Variant calling was carried out using Samtools, Dindel and Pindel^66–69^, and we also used BreakDancer to detect any large structural variants with breakpoints within the exome^68^ (Supplementary Table 6).Variants were annotated using the Ensembl Variant Effect Predictor^70^. For the ES cells and the MBVF line we could not use mapping cross information so quality and impact filtering was carried out instead as the final step (Supplementary Table 6). IGV^71^ was used to check for large deletions in the critical region. Candidate variants were first confirmed by Sanger sequencing, then checked for segregation with the phenotype across the colony as well as all backcross mice (Supplementary Table 3). Protein modelling was carried out using Phyre2^13^ to identify a suitable protein model, and Pymol^72^ to view the model and highlight variant residues. All Sanger sequencing was carried out by Source Bioscience and analysed using Gap4^73^. Venn diagrams were generated using the tool at http://bioinformatics.psb.ugent.be/webtools/Venn/.

### Scanning Electron Microscopy (SEM)

The temporal bones were isolated. The inner ears were dissected out and fixed by 2.5% glutaraldehyde in 0.1M sodium cacodylate buffer with 3 mM calcium chloride at room temperature for 3 hours. Cochleae and the vestibular system were finely dissected in PBS. This was followed by further processing using an osmium-thiocarbohydrazide-osmium (OTOTO) method^74^. The samples were dehydrated in increasing concentrations of ethanol, critical-point dried (Bal-Tec CPD030), mounted and examined under a HITACHI S-4800 or a JEOL JSM 7800F Prime Schottky field emission scanning electron microscope (for *Pcdh^jigl^* and *Espn^spdz^* mice respectively). Images of the organ of Corti were taken at roughly 20% intervals along the cochlear duct and the macula of the utricle and saccule and crista of the ampullae were also imaged. Whole images were adjusted in Photoshop to normalise dynamic range across all panels.

### Middle ear dissection, inner ear clearing and microCT scanning

After culling, the ear canals were checked for cerumen, then the mouse was decapitated and the bulla, tympanic membrane and middle ear cavity inspected for any abnormalities, including the presence of fluid, inflammation or any other obstructions in the middle ear. Observations were recorded on a standard tick sheet. Ossicles were dissected out and stored in 10% formalin. The inner ear was removed and fixed in Bodian’s fixative (75% ethanol, 5% acetic acid, 5% formalin, in water), washed in water and 70% ethanol, then cleared by gentle rotation in 3% KOH for 3 days, changed daily. Samples were then placed in G:E:B (glycerol, 70% ethanol and benzyl alcohol, mixed in a ratio of 2:2:1 by volume) for the final stage of clearing, then stored in G:E (glycerol and 70% ethanol in equal volumes). Images of ossicles and middle ears were taken using a Leica stereomicroscope with a Leica DFC490 camera. To carry out microCT scans, cochleae were immobilized using cotton gauze and scanned with a Scanco microCT 50 to produce 14μm voxel size volumes, using an X-ray tube voltage of 80kVp and a tube current of 80μA. An aluminium filter (0.05mm) was used to adjust the energy distribution of the X-ray source. To ensure scan consistency, a calibration phantom of known geometry (a dense cylinder) was positioned within the field of acquisition for each scan. Test reconstructions on this object were carried out to determine the optimum conditions for reconstruction, ensuring consistency in image quality, and minimising blurring. Reconstruction of the cochlea was performed in Thermo Scientific Amira software.

### Immunohistochemistry and trichrome staining

Samples from adult mice were collected, fixed in 10% formalin, decalcified in 0.1M EDTA, embedded in paraffin wax and cut into 8μm sections. Samples from P4 pups were treated similarly, but no decalcification step was needed. For histological analysis, slides were stained using a trichrome stain, containing Alcian blue, Sirius red and Haematoxylin. Immunohistochemistry was carried out using a Ventana Discovery machine and reagents according to the manufacturer’s instructions (DABMap^TM^ Kit (cat.no 760-124), Haematoxylin (cat.no 760-2021), Bluing reagent (cat.no 760-2037), CC1 (cat.no 950-124), EZPrep (cat.no 950-100), LCS (cat.no 650-010), RiboWash (cat.no 760-105), Reaction Buffer (cat.no 95-300), and RiboCC (cat.no 760-107)). Primary antibodies used were rabbit anti-PMCA2 (Abcam, cat. no: ab3529, diluted 1:500), rabbit anti-MUC13 (Abcam, cat. no: ab124654) and rabbit anti-MYO7A (Proteus, cat. no: PTS-25-6790, diluted 1:100), and the secondary antibody was anti-rabbit (Jackson ImmunoResearch, cat.no 711-065-152, diluted 1:100). Antibodies were diluted in staining solution (10% foetal calf serum, 0.1% Triton, 2% BSA and 0.5% sodium azide in PBS). A Zeiss Axioskop 2 microscope was used to examine slides, and photos were taken using a Zeiss Axiocam camera and the associated Axiocam software. Images were processed in Adobe Photoshop; minimal adjustments were made, including rotation and resizing. Where image settings were altered, the adjustment was applied equally to affected and unaffected samples and to the whole image.

### RNA extraction, RTPCR and qPCR

We collected the brains of adult *Espn^spdz^* (P25) and *rthm* (P28) mice, which were snap-frozen in liquid nitrogen, and the organs of Corti of four-day-old (P4) *Atp2b2^Tkh^* mice, which were dissected out and stored at −20°C in RNAlater stabilisation reagent (Ambion). All RNA dissections were carried out during a fixed time window to avoid circadian variation (*Espn^spdz^*: between 2.5 and 3.5 hours after lights on; *rthm* and *Atp2b2^Tkh^*: between 6 and 7.5 hours after lights on). For the brains, RNA was extracted using TRIzol, in some cases followed by processing through Direct-zol minipreps (Zymo Research, cat. no R2050). For the organs of Corti, RNA was extracted using either QIAshredder columns (QIAgen, cat. no. 79654) and the RNeasy mini kit (QIAgen, cat. no. 74104), or the Lexogen SPLIT kit (Lexogen, cat. no. 008.48), following the manufacturer’s instructions. RNA concentration was measured using a Nanodrop spectrophotometer (ND-8000). RNA was normalised to the same concentration within each litter, then treated with DNAse 1 (Sigma, cat. no: AMPD1) before cDNA creation. cDNA was made using Superscript II Reverse Transcriptase (Invitrogen, cat. no: 11904-018) or Precision Reverse Transcription Premix (PrimerDesign, cat. no: RT-premix2). Primers for sequencing cDNA for testing *Espn* splicing in *Espn^spdz^* mice were designed using Primer3^75^ (Supplementary Table 3). Quantitative RT-PCR on cDNA from *rthm* and *Tkh* mice was carried out on a CFX Connect qPCR machine (Bio-Rad), using probes from Applied Biosystems (Hprt, cat. no: Mm01318747_g1; Jag1, cat. no: Mm01270190_m1; Atp2b2, cat. no: Mm01184578_m1; Slc12a2, cat no: Mm00436563_m1) and Sso-Advanced Master Mix (Bio-Rad, cat. no: 1725281). Relative expression levels were calculated using the 2^-ΔΔct^ equation^76^, with *Hprt* as an internal control. *Jag1* was used to check for the quantity of sensory tissue present in the organ of Corti samples because it is expressed in supporting cells^77, 78^. At least three technical replicates of each sample were carried out for each reaction. The Wilcoxon rank sum test (Mann-Whitney U test) was used to calculate significance for comparisons between *rthm* wildtypes and homozygotes. For the *Atp2b2^Tkh^* qPCR, which involved three groups (wildtype, heterozygote and homozygote), a one-way ANOVA was carried out using SPSS (IBM).

## Acknowledgements

We are grateful to Cassandra Whelan for assistance with the MYO7A staining on the *Atp2b2^Tkh^* and *Tbx1^ttch^* mutants, Seham Ebrahim for additional analysis on the *Atp2b2^Tkh^* mutants, Elysia James for protein modelling of KLHL18, Maria Lachgar-Ruiz for assistance with genotyping, Samoela Rexhaj for help with the mapping of the *rhme* mutation, Hannah Thompson for initial analysis of the *Tbx1^ttch^* mutants, Zahra Hance for her work on the *vthr* mutant, Rosalind Lacey and James Bussell for assistance with mouse colony management, and the Mouse Genetics Project for initial phenotyping of all these mutants. Scanning electron microscropy was carried out at the Wellcome Sanger Institute and King’s College London Centre for Ultrastructural Imaging.

This work was supported by Wellcome (098051, 100669), Medical Research Council (MC_qA137918; G0300212 to KPS.), the European Commission (EUMODIC contract No. LSHG-CT-2006-037188 to KPS), the BBSRC (BB/M02069X/1 to KPS) and King’s College London. AST is funded by the Wellcome Trust (102889/Z/13/Z). TK was supported by the Medical Research Council (MR/L007428/1) and BBSRC (BB/M000281/1). AW was supported by a grant from the Research Foundation Flanders (1700317N). The funders had no role in study design, data collection and analysis, decision to publish, or preparation of the manuscript.

The authors declare no competing interests.

## Author contributions

Electrophysiology was carried out by NJI, MAL, SP, VR and JP. The mapping and segregation testing was done by MAL, JC, SP, RA and MD, and MAL aligned and analysed the exome sequencing data, and carried out the resequencing. Gross morphological analyses of the middle and inner ear were performed by MAL, AW, SR and MD, and scanning electron microscopy was carried out and analysed by JC, FDD and MD. MAL, FDD, RA, and AW contributed to the immunohistochemistry and qPCR. AST reconstructed and analysed the microCT scans. TK and DA carried out exome sequencing and quality control. MAL, NJI, JC, AST, JKW and KPS supervised the research. MAL, NJI, JC and KPS wrote the paper. All authors edited and approved the final manuscript.

## Supplementary Figures and Tables

**Supplementary Fig. 1.**
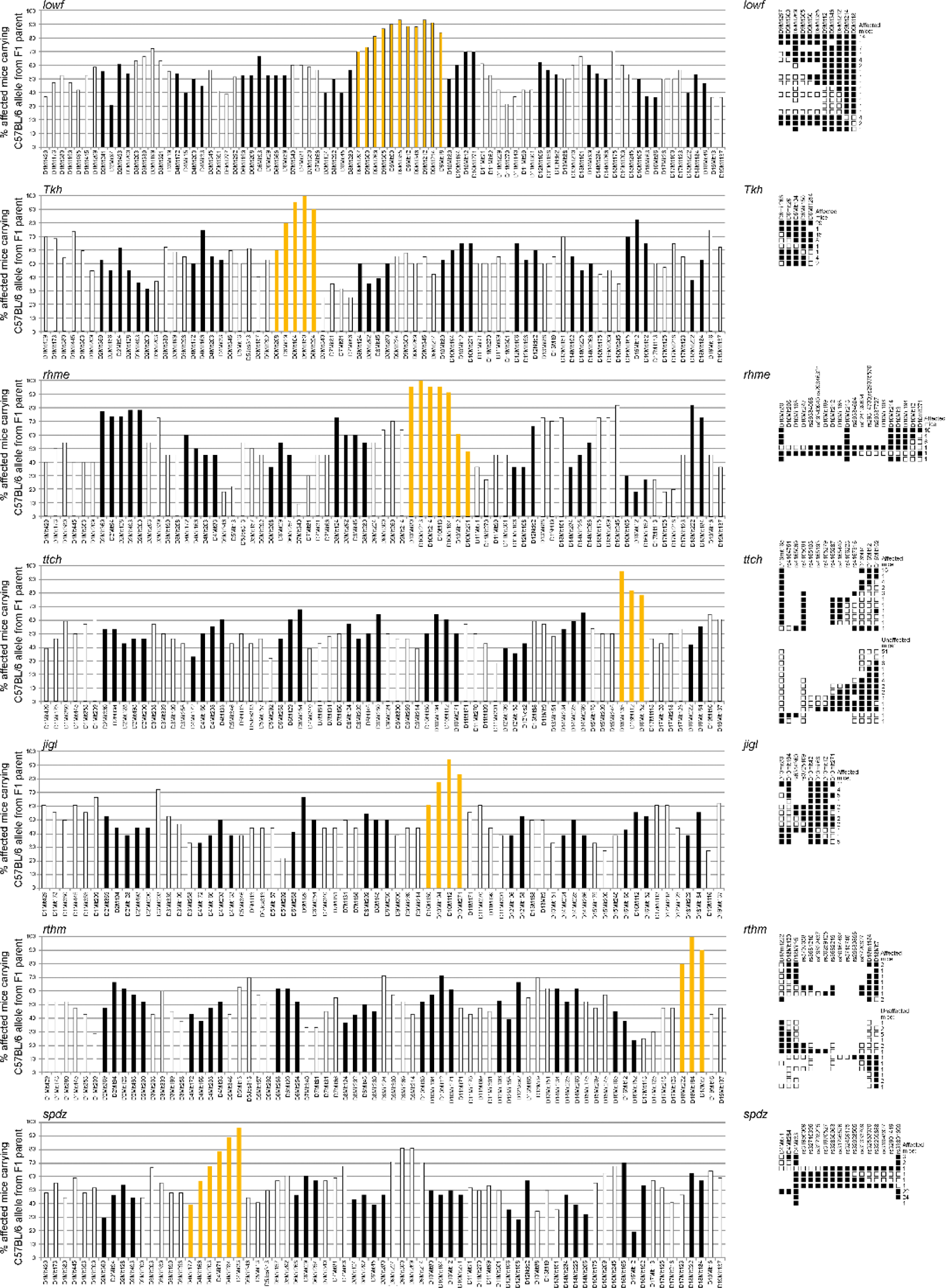
Genome scans and fine mapping. The bar charts show the percentage of affected mice carrying the C57BL/6N allele on the chromosome derived from the outcrossed parent at each marker. Alternate chromosomes are indicated by different shading, and the linked chromosome for each mutation is coloured gold. Following the initial mapping, the critical region was identified using more markers and, where necessary, SNVs known to differ between C57BL/6N and C3HeB/FeJ, the two strains used for the mapping crosses. On the right, the haplotype charts show the fine mapping of each allele. Each row represents the marker pattern for the chromosome derived from the F1 parent, with white boxes representing the C3HeB/FeJ-like marker types and black boxes representing the C57BL/6N-like marker types. A change from black boxes to white or vice versa indicates that a recombination event has occurred during meiosis leading to the production of that chromosome in the F1 parent. For all alleles except *Tkh*, the F1 parent was backcrossed to the affected parental line to obtain unaffected mice heterozygous for the causative mutation and affected homozygotes, but because the *Tkh* allele was semidominant, the F1 parent was backcrossed to C3HeB/FeJ, resulting in affected mice heterozygous for the causative mutation and unaffected wildtypes. The number to the right of each row indicates the number of mice showing that particular pattern of marker types. Numbers: *lowf*: n=30 affected mice for the genome scan, 44 affected mice for fine mapping; *Tkh*: n = 20 affected mice for the genome scan, 61 affected mice for fine mapping; *rhme*: n = 11 affected mice for the genome scan, 23 affected mice for fine mapping; *ttch*: n = 28 affected mice for the genome scan, 31 affected and 75 unaffected mice for fine mapping; *jigl*: n = 18 affected mice for the genome scan, 35 affected mice for fine mapping; *rthm*: n = 21 affected mice for the genome scan, 9 affected and 19 unaffected mice for fine mapping; *spdz*: n = 28 affected mice for the genome scan, 56 affected mice for fine mapping.

**Supplementary Fig. 2.**
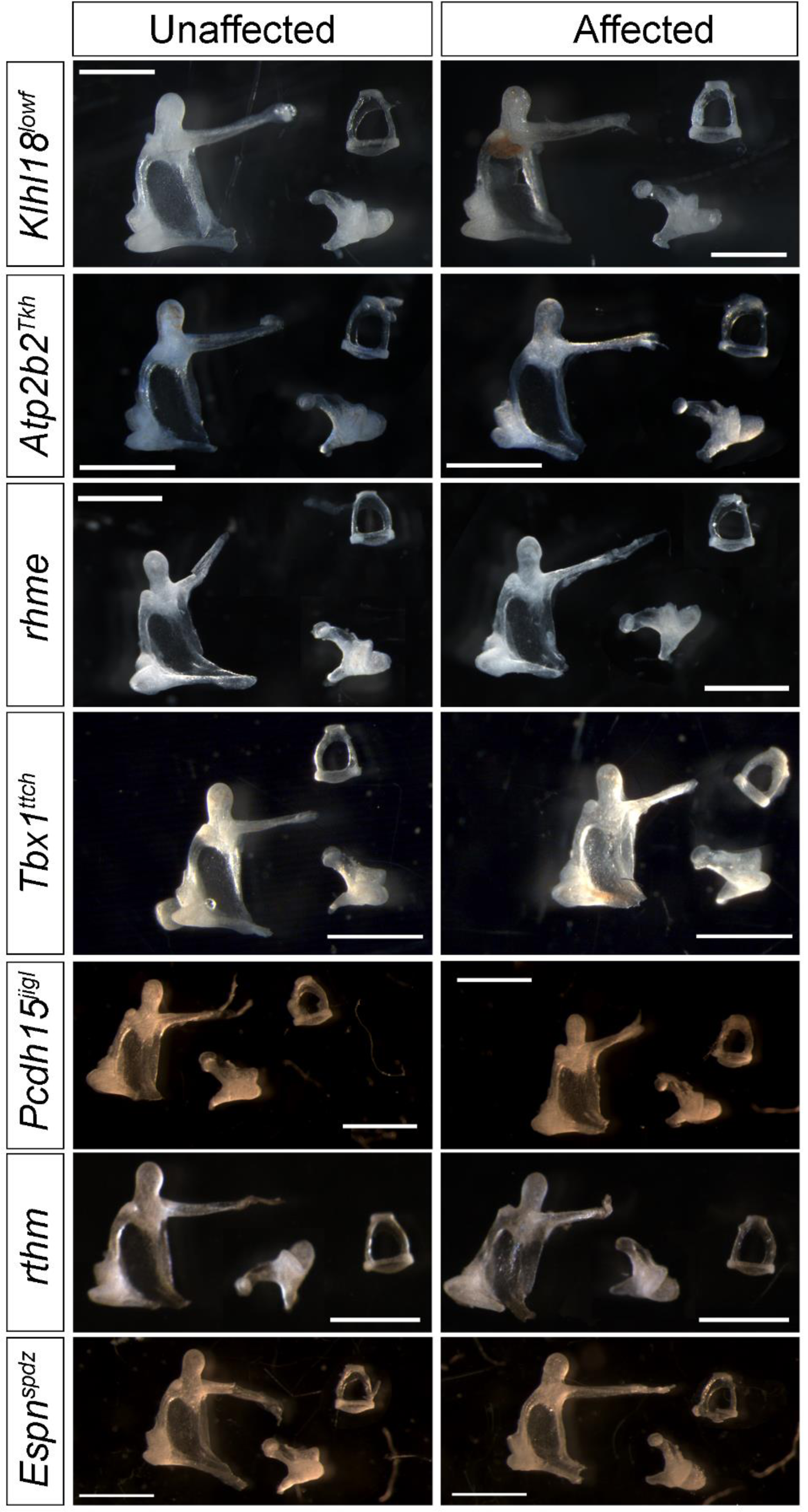
Middle ear ossicles from mice displaying hearing loss show no defects. Ossicles from affected mice from all seven lines (right) and unaffected littermates (left). Numbers: *Klhl18^lowf^* n=5 unaffected, 9 affected mice at P94±1day; *Atp2b2^Tkh^* n=3 unaffected (wildtype), 3 affected (homozygote) mice at P36±2 days; *rhme* n=3 unaffected, 3 affected mice at P69; *Tbx1^ttch^* n=3 affected, 3 unaffected mice at P28 or older; *Pcdh15^jigl^* n=7 affected, 4 unaffected mice at P30±1 day; *rthm* n=3 affected, 3 unaffected mice at P28±1 day; *Espn^spdz^* n=5 affected, 4 unaffected mice at P57 or older. Scale bar = 1mm. For each panel, the malleus is shown on the left, the stapes on the top right and the incus on the bottom right.

**Supplementary Fig. 3.**
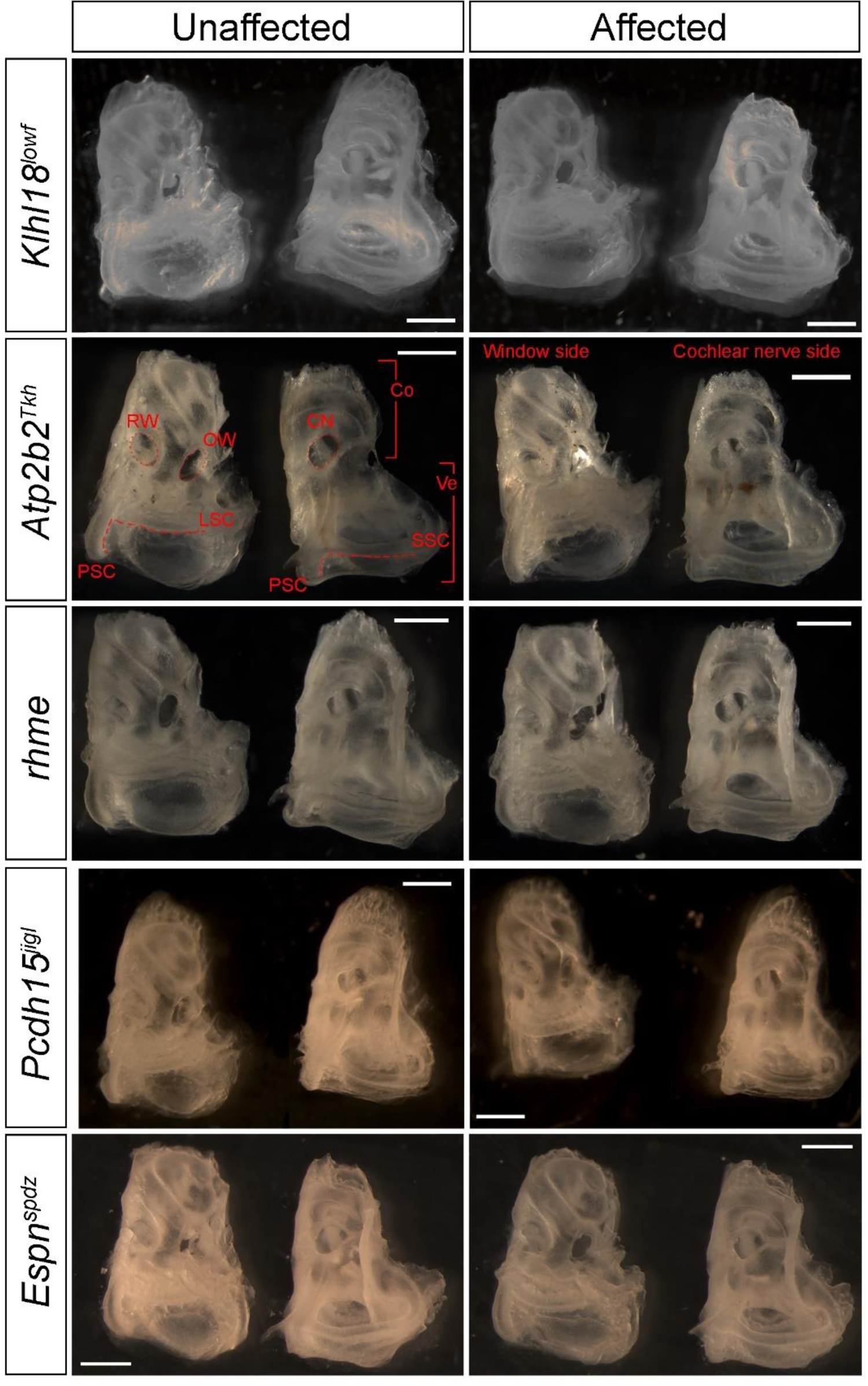
Cleared inner ears from mice displaying hearing loss which show no gross defects. Cleared inner ears from affected mice carrying the *Klhl18^lowf^*, *Atp2b2^Tkh^*, *rhme*, *Pcdh15^jigl^* and *Espn^spdz^* alleles (right) and unaffected littermates (left). Numbers: *Klhl18^lowf^* n=5 unaffected, 7 affected mice at P43±1day; *Atp2b2^Tkh^* n=3 unaffected (wildtype), 3 affected (homozygote) mice at P36±2 days; *rhme* n=3 unaffected, 3 affected mice at P69; *Pcdh15^jigl^* n=12 affected, 8 unaffected mice at P30±1 day; *Espn^spdz^* n=5 affected, 4 unaffected mice at P57 or older. Scale bar = 1mm. Labels have been added to the *Atp2b2^Tkh^* inner ears. The middle ear side, with the round (RW) and oval (OW) windows is shown on the left of each panel, and the brain side, where the cochlear nerve exits (CN), on the right. The round and oval windows and the cochlear nerve exits are marked by dotted lines, and the semicircular canals by dashed lines. Brackets indicate the cochlea (Co) and vestibular region (Ve). LSC=lateral semicircular canal; SSC=superior semicircular canal; PSC=posterior semicircular canal.

**Supplementary Fig. 4.**
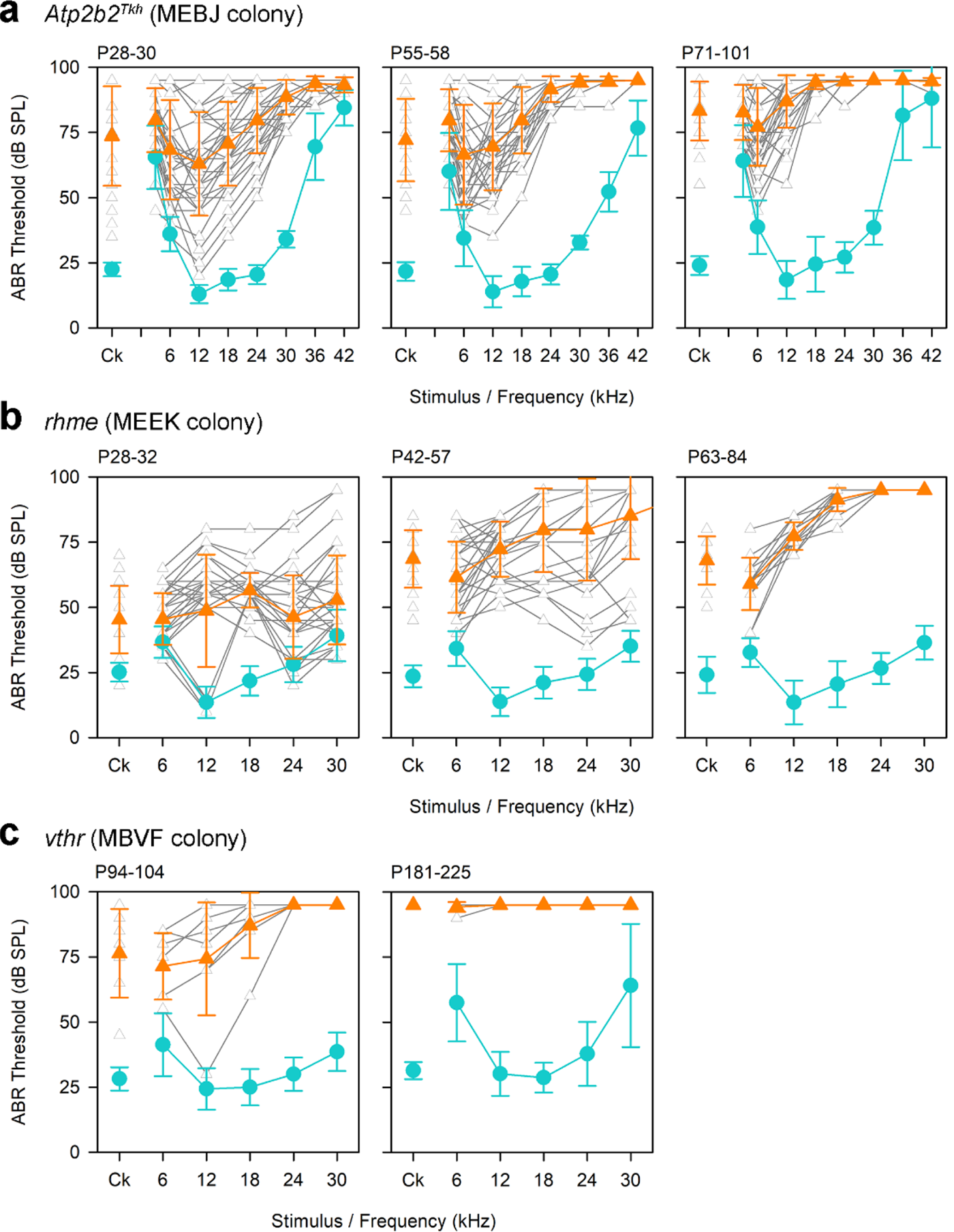
ABR thresholds of mice from the MEBJ, MEEK and MBVF lines showing progressive division into affected and unaffected mice by threshold. This figure shows ABR thresholds from mice from the MEBJ (a), MEEK (b) and MBVF (c) colonies, showing both the progression of hearing loss with age and the phenotypic division into affected and unaffected mice. These graphs include both genotyped mice (which will also be represented in other figures) and those mice tested before genotyping was possible. Where possible, the same mice were tested at each age. **a** ABR thresholds of affected MEBJ mice carrying the *Atp2b2^Tkh^* allele and unaffected littermates, showing the progression of hearing loss in affected mice (orange triangles) compared to unaffected mice (teal circles) (n = 10 unaffected and 37 affected at P28-30; n=9 unaffected and 31 affected at P55-58; n=19 unaffected and 37 affected at P71-101). Individual audiograms from affected mice are shown in grey. **b** ABR thresholds from affected (orange triangles) and unaffected (teal circles) *rhme* (MEEK) mice at P28-32 (n = 42 unaffected, 35 affected), P42-57 (n = 83 unaffected, 28 affected) and P63-84 (n = 28 unaffected, 15 affected). Traces from individual affected mice are shown in grey. **c** ABR thresholds of seven MBVF mice carrying the unknown *vthr* mutation at P94-104 and unaffected mice from the same colony (n=7 affected, 89 unaffected) and P181-225 (n=7 affected, 28 unaffected). The hearing loss appears to be variable at the younger age but progresses to be more severe at ages over 6 months. Traces from individual affected mice are shown in grey.

**Supplementary Fig. 5.**
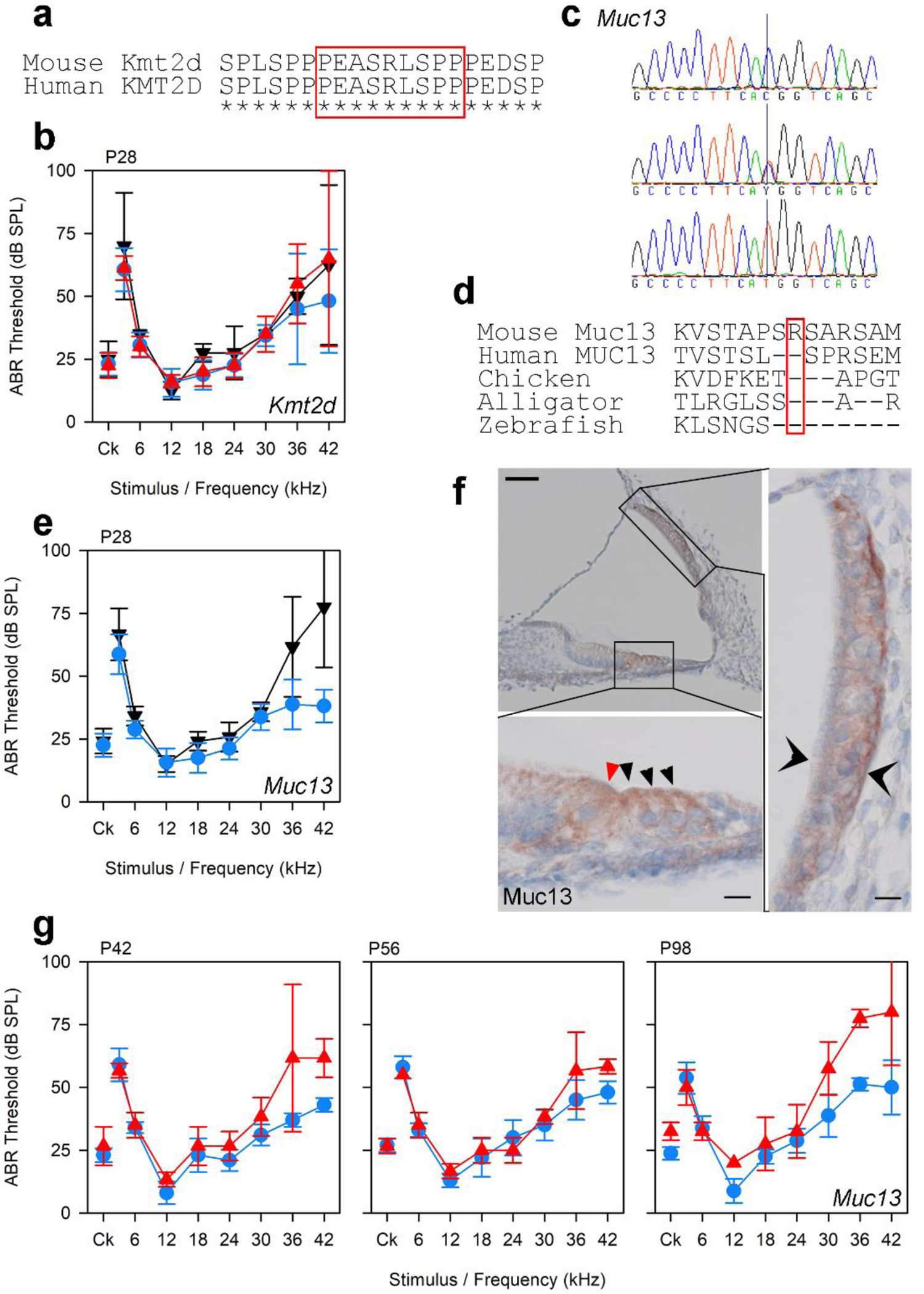
Two other mutations found in the MDLY line carrying the *ttch* allele did not affect hearing. **a** Clustal alignment showing deleted amino acids in *Kmt2d* (red box) caused by the g.15:98863687_98863713del mutation, resulting in a loss of 9 amino acids (ENSMUST00000023741, p.586-594del). The region is conserved between mouse and human but not in other vertebrates checked. **b** ABR thresholds for P28 mice homozygous (n=4, red triangles), heterozygous (n=8, blue circles) or wildtype (n=2, black inverted triangles) for the *Kmt2d* deletion. All these mice were heterozygous or wildtype for the *Tbx1^ttch^* allele, and mice homozygous for the deletion have no difference in their ABR thresholds. **c** Sequence trace showing the missense mutation in *Muc13* (g.16:33807881C>T). **d** Clustal alignment showing the affected amino acid in *Muc13* (red box; p.(Arg334Trp), ENSMUST00000115044). It is not conserved in any other vertebrates checked. **e** ABR thresholds for P28 mice heterozygous (n=8, blue circles) or wildtype (n=6, black inverted triangles) for the *Muc13* missense mutation. All these mice were heterozygous or wildtype for the *Tbx1^ttch^* allele. **f** MUC13 expression at P4 in a mouse heterozygous for both the *Muc13* missense variant and the *Tbx1^ttch^* allele (n=2). Brown indicates the presence of MUC13 protein, which is visible in the hair cells (arrowheads; red points to the inner hair cell and black to the outer hair cells) and in the basal cells of the stria vascularis (twin open arrowheads). Scale bar = 50µm in the top left panel and 10µm in the two high magnification panels. The section shown is from the region 50% of the distance along the organ of Corti from base to apex. **g** ABR thresholds for mice homozygous (red triangles) or heterozygous (blue circles) for the *Muc13* mutation at later ages (P42 and P56: n=5 heterozygotes, 3 homozygotes; P98 n=4 heterozygotes, 2 homozygotes). All these mice were heterozygous or wildtype for the *Tbx1^ttch^* allele.

**Supplementary Fig. 6.**
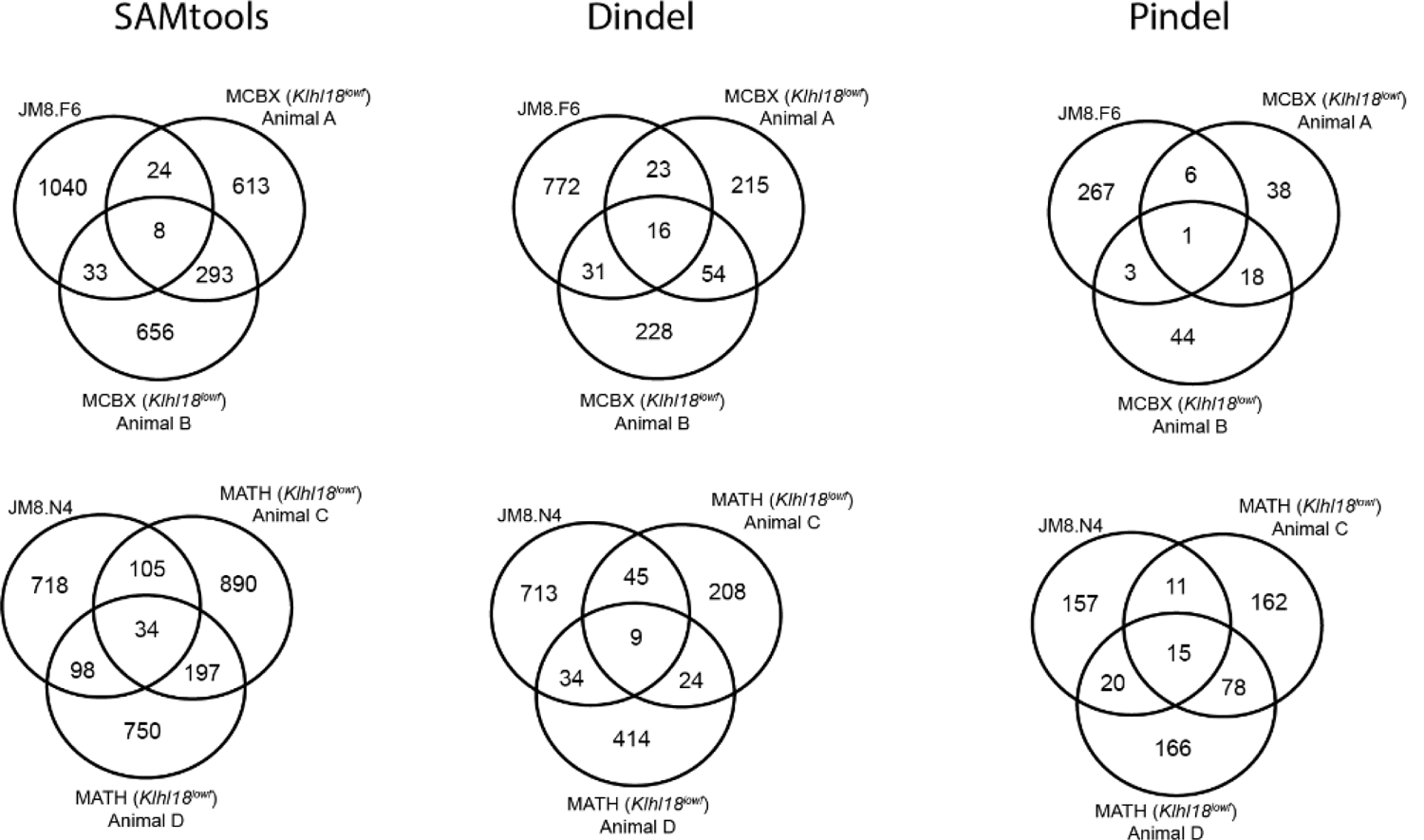
A subset of variants identified by each caller were passed on from parental ES cells to mice created using that ES cell line. Venn diagrams showing the number of high quality variants in two ES cell lines (JM8.F6 and JM8.N4) which were shared by mice from lines created using those ES cell lines (MCBX and MATH), detected by SAMtools, Dindel and Pindel. Both these mouse lines also carry the spontaneous *Klhl18^lowf^* mutation, which was not seen in any of the ES cell lines. None of the variants identified by BreakDancer were shared between the ES cells and the mice (Supplementary Table 4).

**Supplementary Fig. 7.**
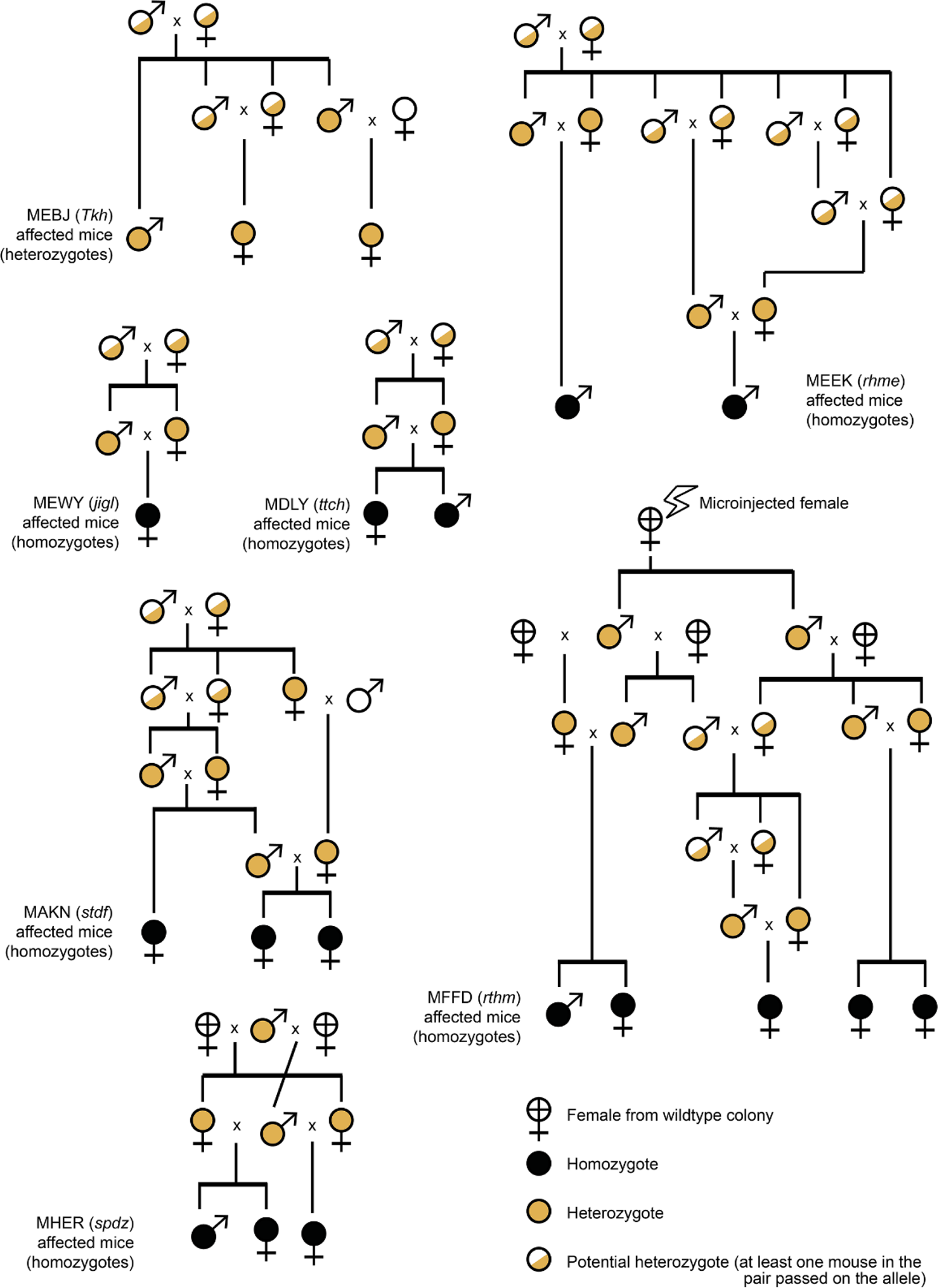
Pedigrees of the affected mice showing the latest possible point at which each mutation could have arisen. Pedigrees of mouse lines for seven of the eight mutations described here. The *Klhl18^lowf^* allele is thought to have arisen in the wildtype colony and is not shown. For each of the other lines, the pedigree shows the latest possible point at which the mutation could have arisen. Homozygotes are shown in solid black circles, heterozygotes (obligate carriers for the recessive mutations or mice carrying the semidominant *Atp2b2^Tkh^* allele) are shown in yellow, and where either or both mice in a pair may have been carrying the mutant allele, this is indicated by the white/yellow colour. In only one line (MFFD, *rthm*) does the mutation definitely originate at or before the point of microinjection of the targeted ES cells. Only the mice at the bottom of each pedigree, marked by the text, are known to have been affected (and therefore homozygotes or, for *Atp2b2^Tkh^*, heterozygotes); all the other genotypes and phenotypes are inferred.

**Supplementary Table 1.**
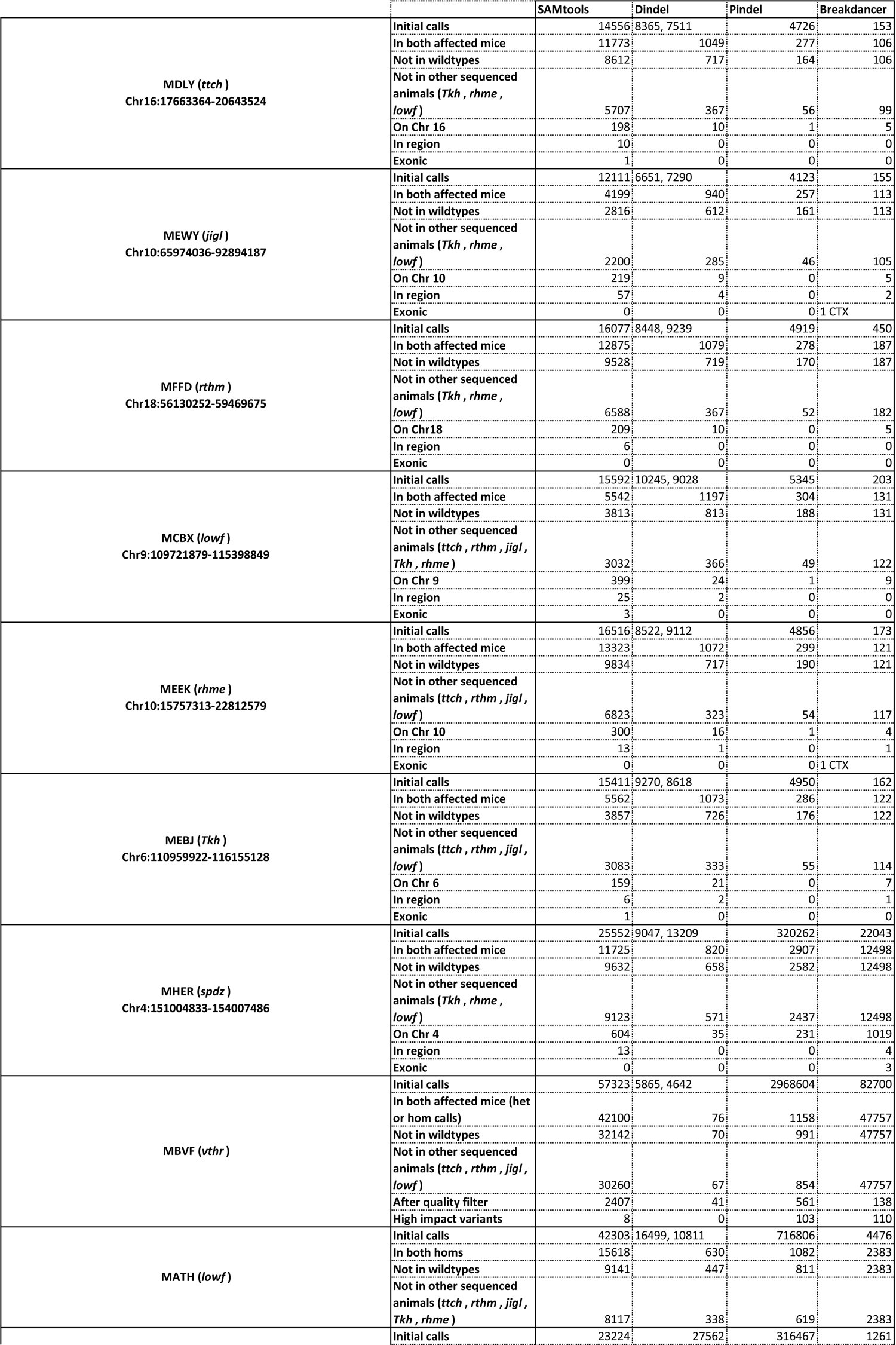
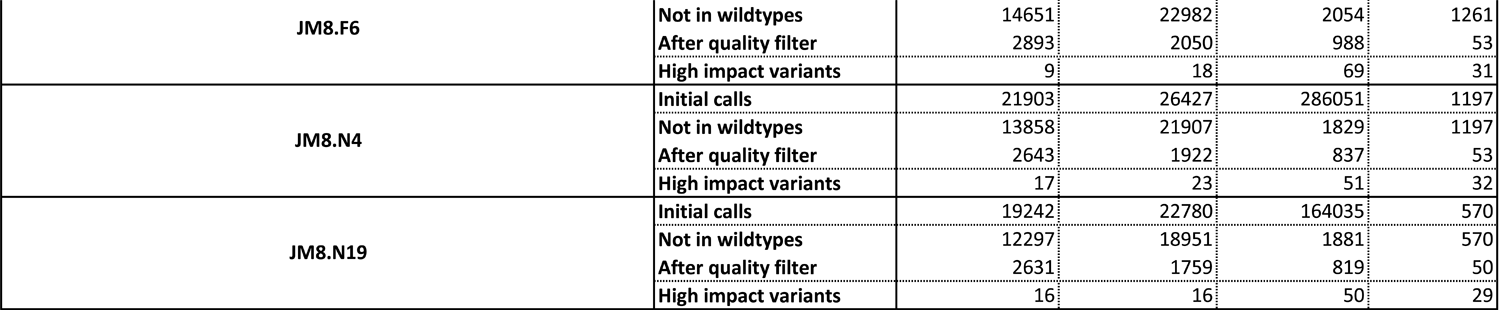
Variant counts from whole exome sequencing. This table shows the variant counts from each caller for all mice and ES cell line sequenced, and the filtering steps undertaken (the identification of the *S1pr2 ^stdf^* mutation has been previously described^12^). The lefthand column holds the line name, with the allele in brackets where applicable, and the nonrecombinant region identified from mapping, where that was carried out. SAMtools, Pindel and BreakDancer call variants from both animals sequenced, so the single number represents all variants seen in one or both mice. Dindel calls indicate variants per animal. For the “not in wildtype” filter, variants were compared to known variants from the Ensembl database and the Mouse Genomes Project^63, 82, 83^, and only novel variants were kept. We also carried out an additional filtering step, removing any variants seen in the other mice sequenced in this project which had markedly different phenotypes (since the MHER (*spdz*) line was sequenced much later than the others, variants from this line were not used in this filter). For two of the lines, MATH and MBVF (*vthr*), we were not able to map the mutation. In the case of the MATH line, the affected mice displayed the same low frequency hearing loss phenotype as the MCBX line, and we found the same mutation (*Klhl18 ^lowf^*) in all the affected mice we were able to sequence, so we did not filter variants any further. For the ES cells and the *vthr* mice (MBVF colony), we carried out two additional filters; a quality filter (Supplementary Table 6) and an impact filter, where only variants predicted to have a high impact on gene function (by the Ensembl Variant Effect Predictor) passed the filter.

**Supplementary Table 2.**
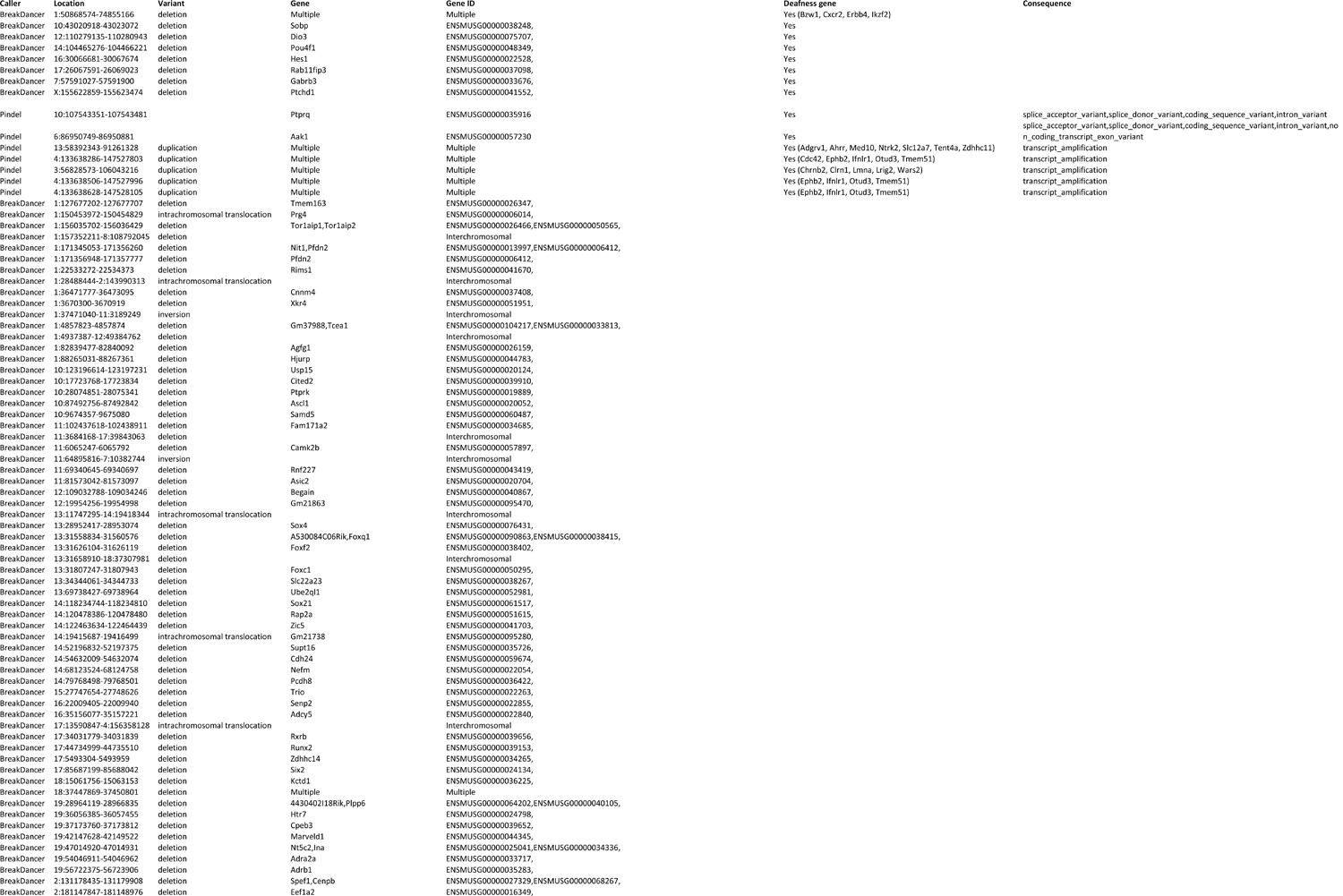
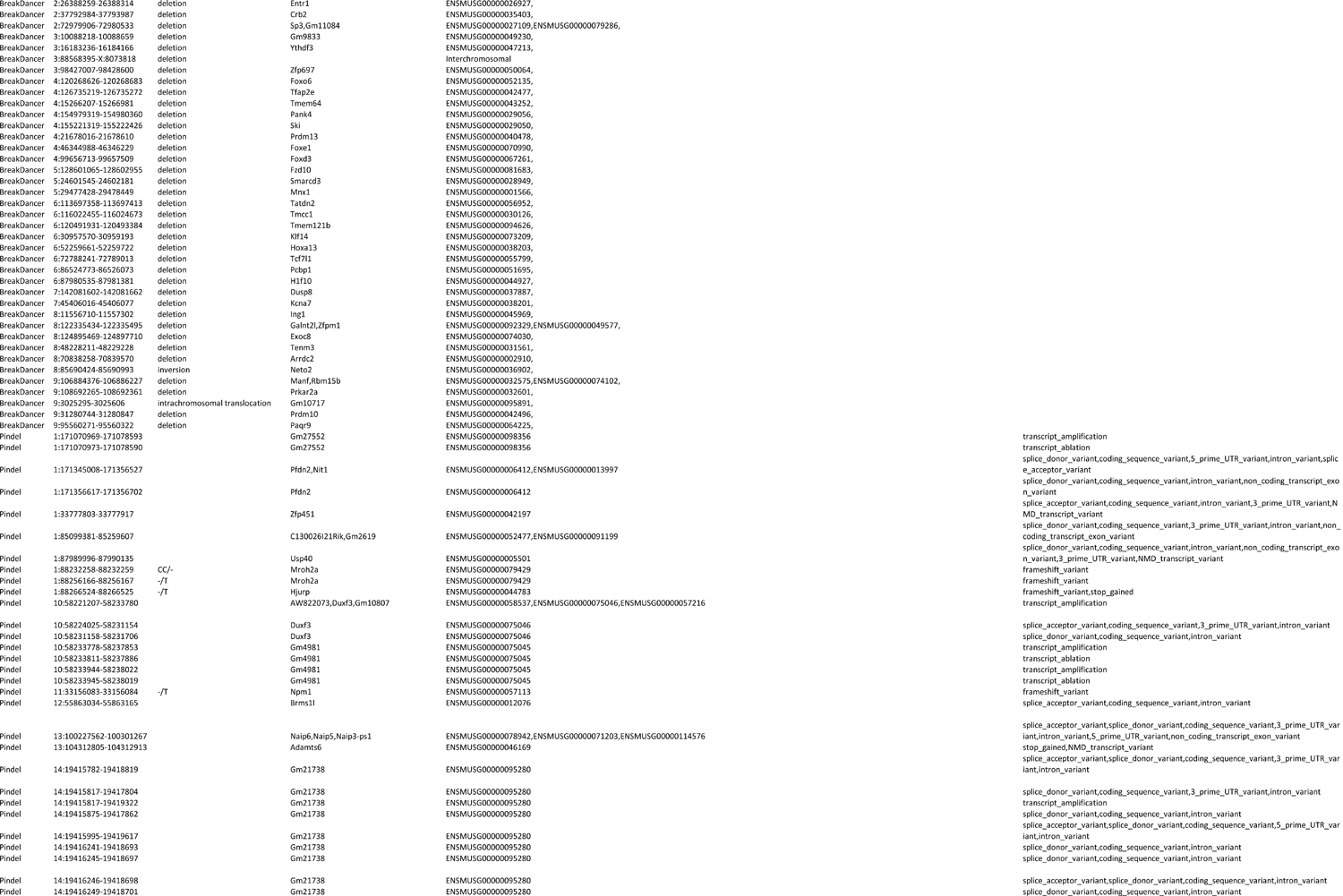
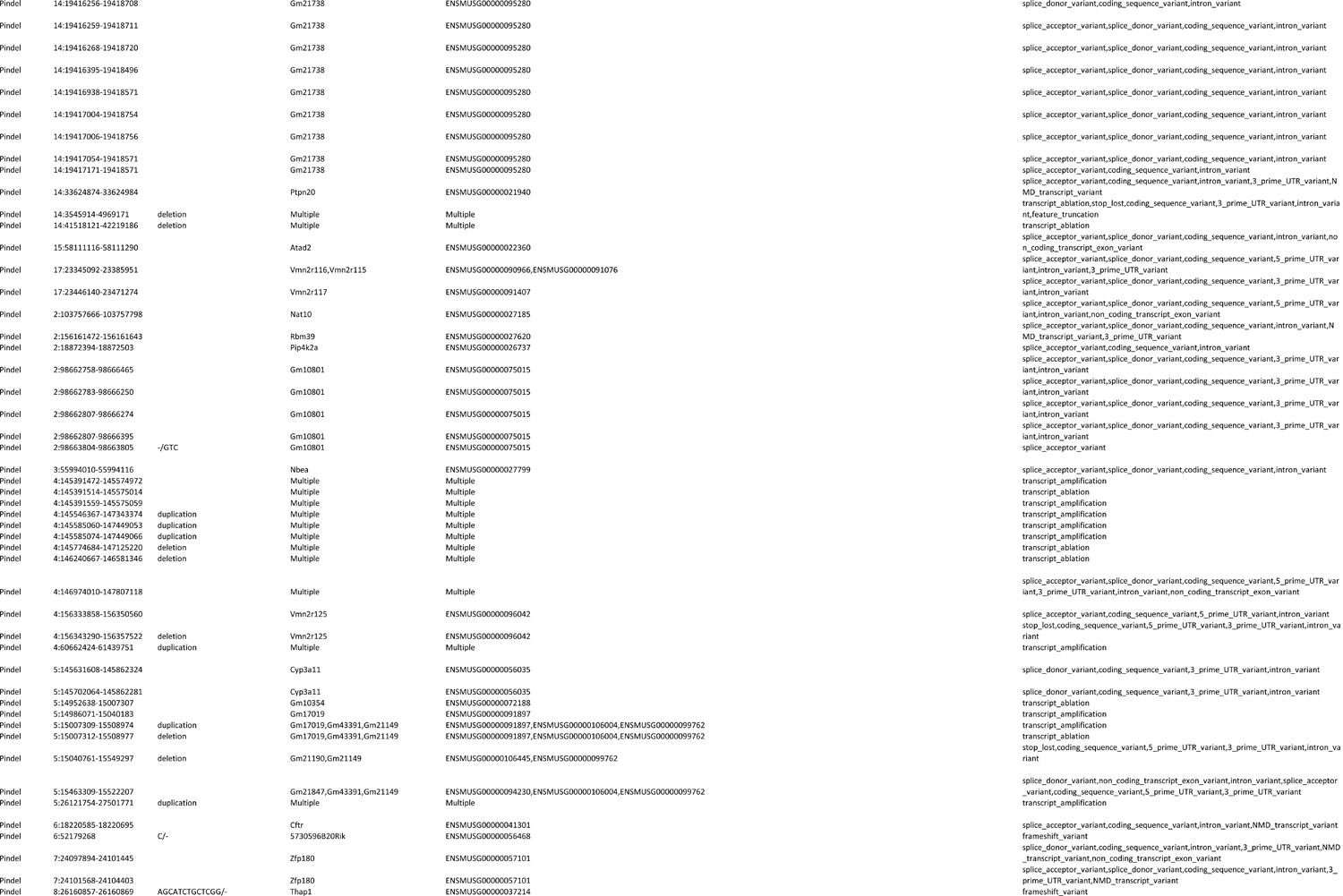
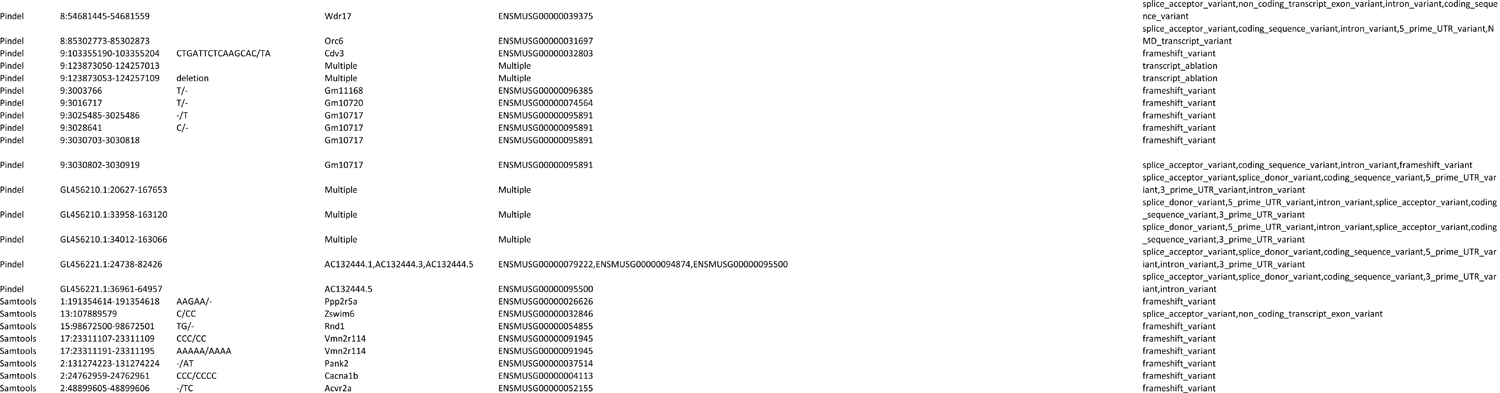
High quality, high impact variants identified in *vthr* mice (MBVF colony)

**Supplementary Table 3.**
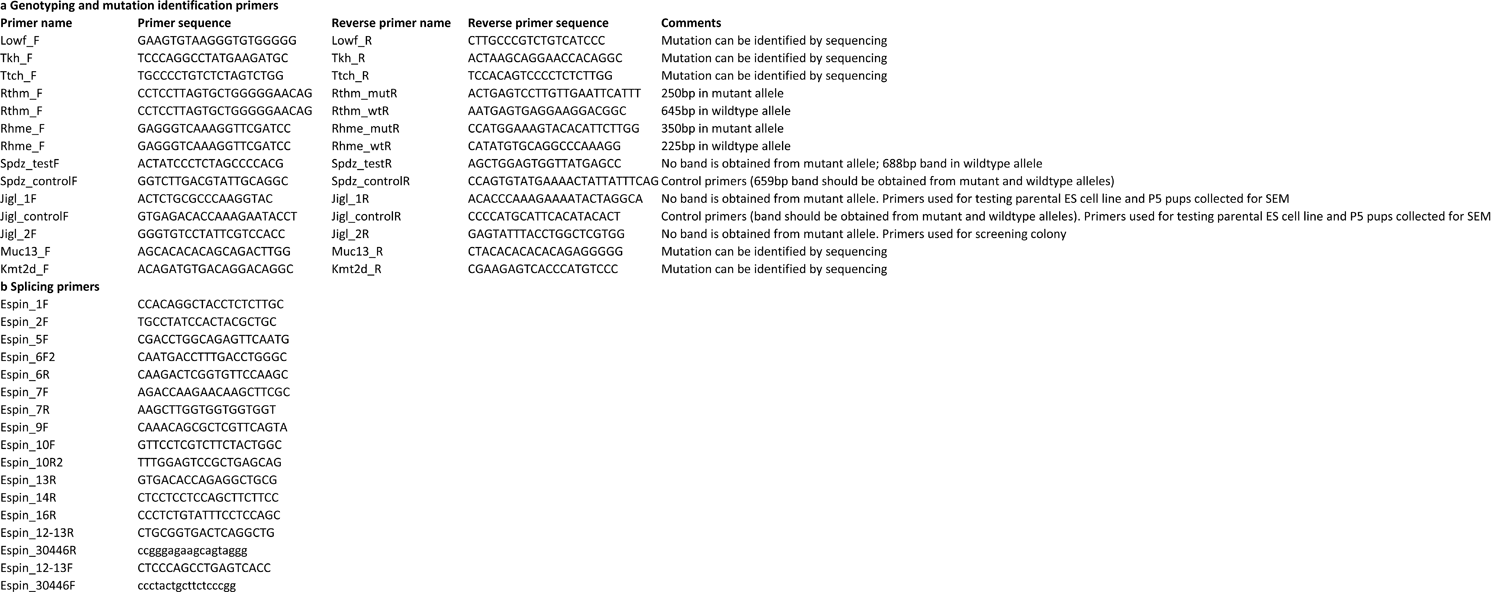
Primer sequences for primers designed in this study. **a** Primers used for genotyping spontaneous alleles. **b** Primers used to examine *Espn* mRNA splicing in mice carrying the *spdz* allele.

**Supplementary Table 4.**
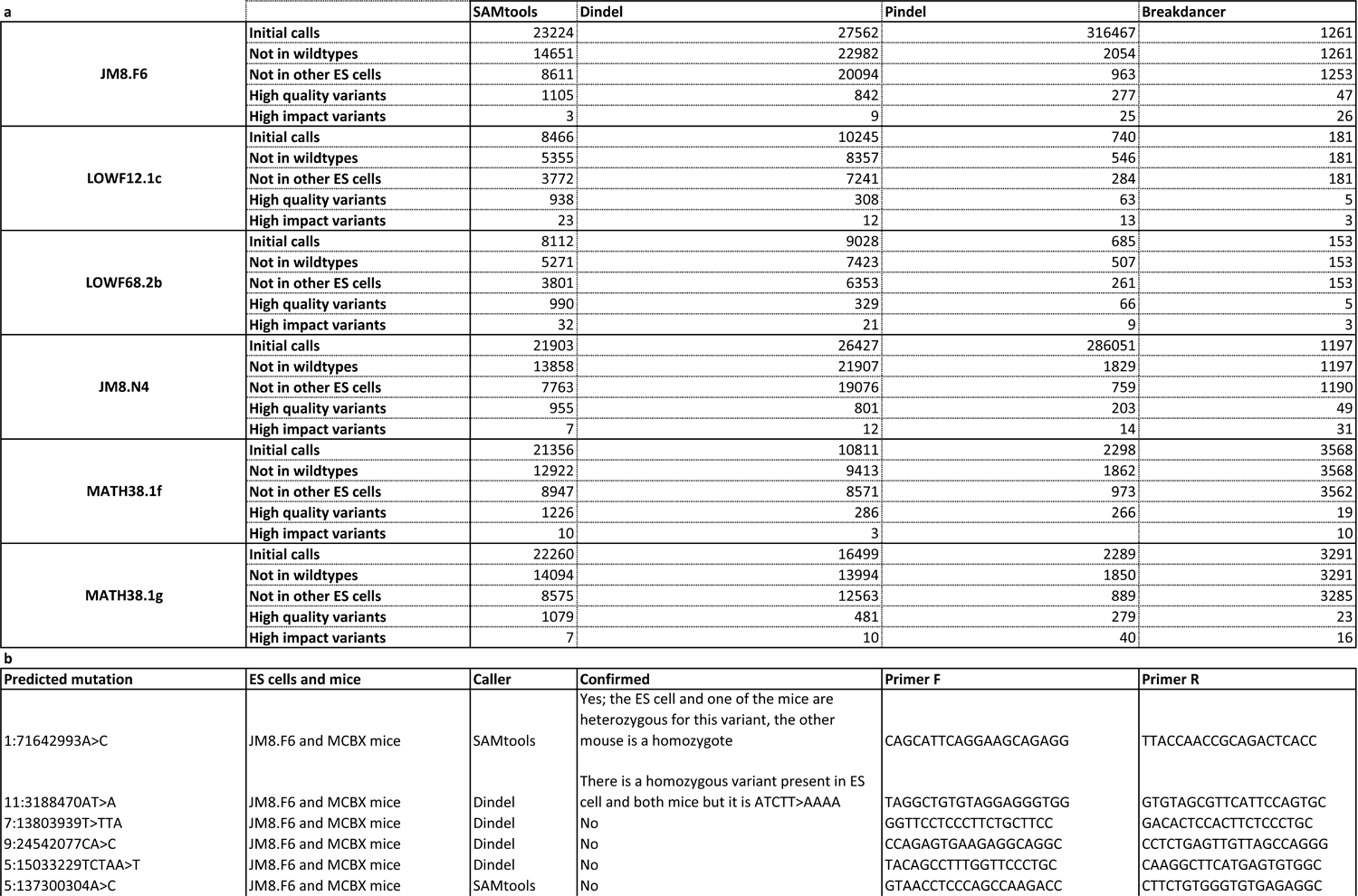
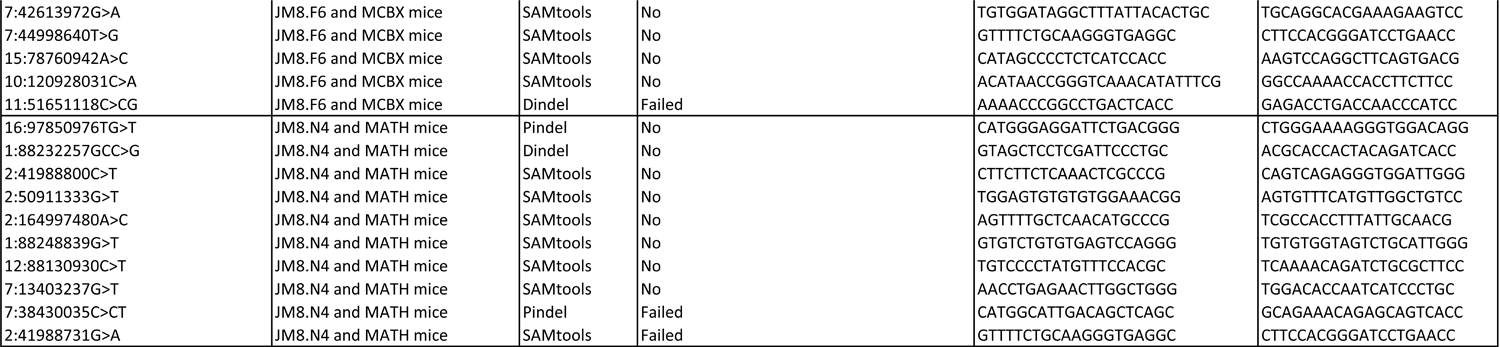
ES cell variant counts and confirmation. **a** This table shows the variant counts from each caller for individual mice and their ancestral ES cells, and the filtering steps undertaken. The process is similar to that described for Supplementary Table 1, but here instead of filtering by variants found in other mouse lines, we eliminated variants found in other ES cells. The quality filter steps are defined in Supplementary Table 6, and high impact variants were those predicted by the Variant Effect Predictor to have a high impact on gene function. **b** Variant validation using Sanger sequencing and primers used. Most variants were not validated. Three of them failed (either by failure to amplify or failure to obtain good quality sequencing), and two were confirmed in the JM8.F6 ES cell line and its descendant mice, although one of them, the small indel called by Dindel, was misidentified.

**Supplementary Table 5.**
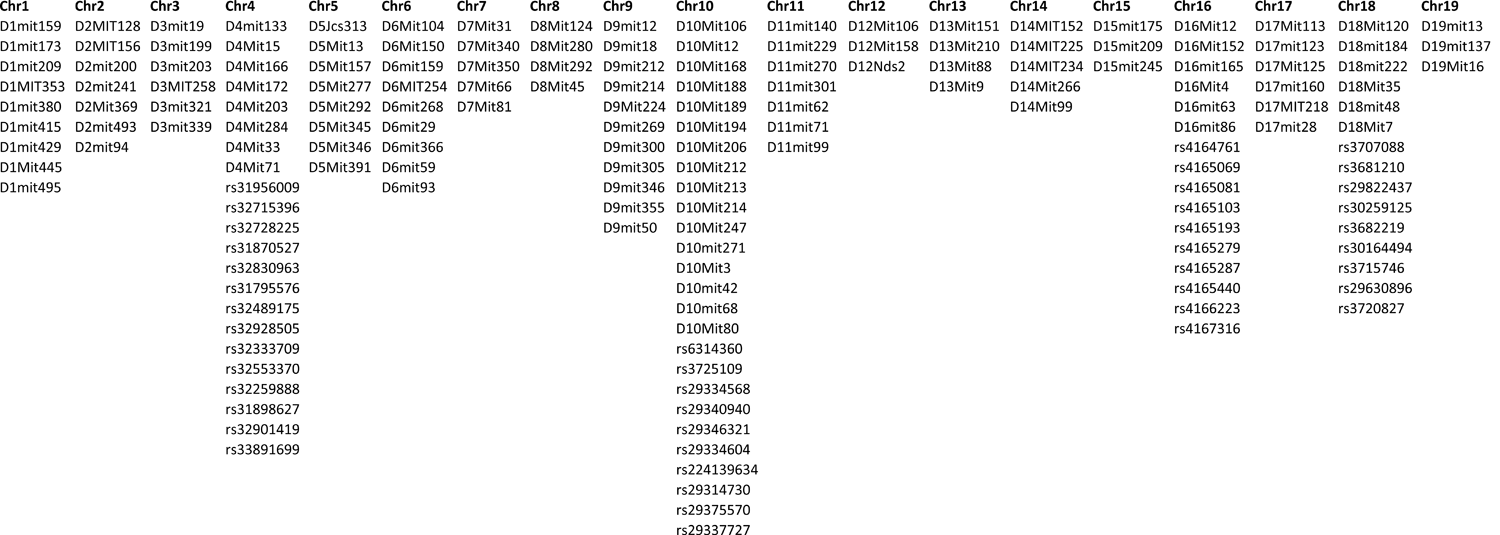
Markers used for genome scans and fine mapping.

**Supplementary Table 6.**
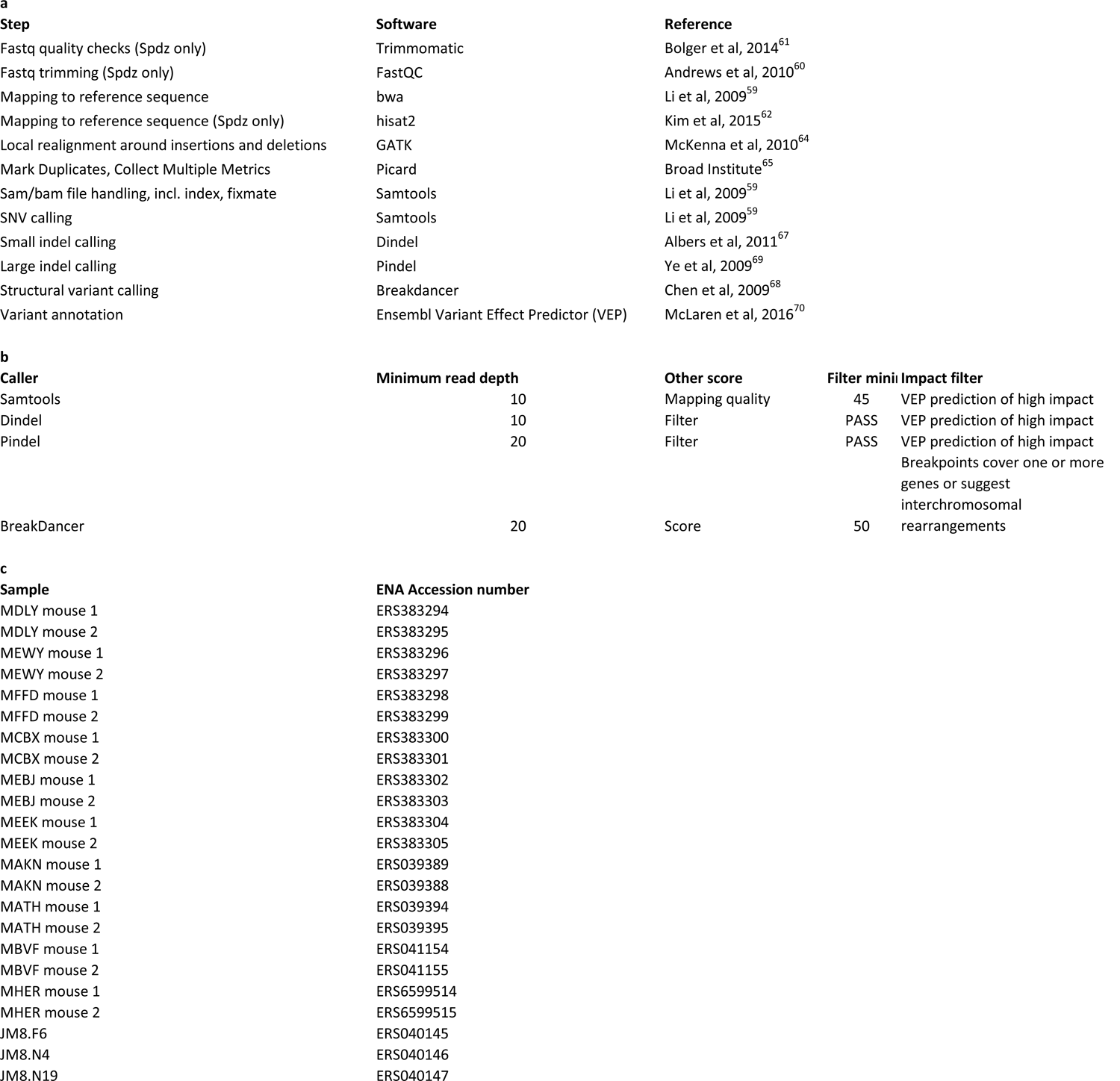
Sequence processing and filtering details. **a** Software used for processing and aligning reads, and calling and annotating variants. **b** Thresholds chosen for the variant quality filtering carried out for the MBVF line and ES cell whole exome sequencing. **c** ENA accession numbers for individual samples.

